# Characterization of alternative mRNA splicing associated with tumor thrombus in clear-cell renal cell carcinoma

**DOI:** 10.1101/2024.09.26.615158

**Authors:** Zeev Cohen, Harel Reinus, Iddo Ben-Dov, Tomer Kalisky

## Abstract

Approximately 10% of Renal Cell Carcinoma (RCC) tumors form a thrombus that invades into nearby vasculature and is associated with lower survivability. However, the mechanisms driving tumor thrombus formation and progression are poorly understood. We therefore examined a publicly available RNAseq dataset of samples from RCC patients containing tumors with and without thrombi, as well as associated normal kidney tissues, and compared them. Using cell deconvolution, we found indications that kidney tumors and thrombi are associated with loss of mature kidney-specific cell types and gain of proliferative characteristics. Moreover, we identified a set of transcripts that are alternatively spliced between tumors with and without thrombi. Using motif enrichment analysis for known RNA binding proteins we found putative splicing regulators that are presumably associated with thrombus formation. We believe that this study will assist in unraveling cellular and molecular mechanisms of tumor progression and thrombus formation in RCC.

## INTRODUCTION

Renal cell carcinoma (RCC) accounts for approximately 4% of all adult malignancies. The most common subtype is clear-cell RCC (ccRCC) (Almatari et al. 2023; Pallagani et al. 2021), which is thought to originate from the proximal tubule of the adult kidney (Muglia and Prando 2015). An estimated 4-10% of patients with RCC will develop a tumor thrombus (TT) (Mootha et al. 1999; Quencer et al. 2017; Topaktaş et al. 2019), an extension of the tumor into surrounding blood vessels that is associated with poor prognosis and low survival. Tumor thrombi typically invade and occlude the inferior vena cava (IVC), and in extreme cases can also reach the heart, thus complicating tumor resection surgery (Almatari et al. 2023; Mootha et al. 1999; Topaktaş et al. 2019).

The cellular and molecular mechanisms behind tumor thrombus formation in RCC, and their relation to other metastatic processes, are poorly understood. Wang et al. (X.-M. Wang et al. 2020) sequenced RNA of ccRCC samples collected from 152 Chinese patients. They found differences between tumors with and without thrombi in genes related to immune cell composition, angiogenesis, cell division, and the epithelial to mesenchymal transition (EMT). Shi et al. (Shi et al. 2022) used single cell RNAseq to compare samples from thrombi and matched primary tumors from 8 ccRCC patients, and found that thrombi are enriched with tissue resident CD8+ T cells in a progenitor exhausted state, have enhanced extracellular matrix (ECM) remodeling activity, and have a lower expression of immunosuppressive markers. Kim et al. (K. Kim et al. 2021) used histology and bulk RNA-Seq to compare thrombi with their associated primary tumors from 82 ethnically-diverse patients, and found that the thrombi had higher expression of genes that promote the EMT such as PRRX1. They also over-expressed immediate-early genes such as the transcription factors EGR1 and EGR3, as well as FOSB, FOS, and JUNB, that encode components of the AP-1 transcription factor complex. Moreover, they found that grade of the tumor thrombus is a predictor of metastasis of the primary tumor.

There is evidence that alternative mRNA splicing may also be involved in tumor formation and progression (Ouyang et al. 2021; Pradella et al. 2017). Alternative mRNA splicing enables a single gene to encode multiple transcripts, each possibly having a different function, thus allowing the same gene to assume multiple roles during tissue formation and regeneration. It was previously found that alternative splicing in the genes FGFR2 (Hovhannisyan, Warzecha, and Carstens 2006; Ranieri et al. 2018; Warzecha, Sato, et al. 2009), CD44 (Brown et al. 2011; Vos et al. 2016), ENAH, and CTNND1 (Warzecha, Sato, et al. 2009; Warzecha, Shen, et al. 2009) is associated with tumor progression and EMT. Therefore, in this study we set to characterize alternative mRNA splicing associated with tumor thrombi in ccRCC. We first downloaded a publicly available RNAseq dataset of samples collected from RCC patients containing tumors with and without thrombi, as well as associated normal kidney tissues. Then, we identified transcripts that are alternatively spliced between tumors with and without a thrombus. Finally, using motif enrichment analysis, we found putative splicing regulators that are presumably associated with thrombus formation. We believe that understanding the mechanisms of alternative mRNA splicing and their association with tumor formation, progression, and invasion, will lead to development of novel therapeutic strategies (Baughn et al. 2023).

## MATERIALS AND METHODS

### Datasets and Preprocessing

RNA sequencing reads generated by Wang et al. (X.-M. Wang et al. 2020) were downloaded from the NCBI Sequence Read Archive (SRA) (accession code PRJNA596338) using SRA tools. Patient clinical metadata was downloaded from the journal’s website. STAR (version 2.7.10b) (Dobin et al. 2013) was used to align the reads to the human reference genome (hg38) and to obtain a raw counts matrix. DESeq2 (version 1.42.0) (Love, Huber, and Anders 2014) was used to obtain a normalized counts matrix.

### Cell Deconvolution

BisqueRNA (version 1.0.5) (Jew et al. 2020) was used to estimate cell type proportions within each bulk RNA sample. As a reference single-cell RNAseq gene expression dataset, we used the adult human kidney dataset downloaded from the Kidney Cell Atlas (https://www.kidneycellatlas.org/) (Stewart et al. 2019). Since BisqueRNA requires the single-cell reference as a Seurat object, we used the ‘scanpy’ python library (version 1.9.6) to convert the single-cell reference dataset from H5ad format to Market Exchange Format (MEX) that can be read by Seurat (version 5.0.1) (Hao et al. 2024). Heatmaps were generated using the ‘complexHeatmap’ R package (version 2.18.0).

### Alternative splicing

rMATS-turbo (version 4.12) (Y. Wang et al. 2024) was used to detect alternatively spliced transcripts. In each rMATS run we compared two sets of samples chosen according to their position in PCA latent space. Additionally, to obtain the inclusion levels across all samples and all splicing events, we performed a separate rMATS run with the option ‘--cstat 0’ on all samples. Alternative splicing in selected transcripts was visually validated using the Integrated Genome Viewer (IGV) (Robinson et al. 2023). Sashimi plots were generated using the ‘ggsashimi’ command-line tool (version 1.1.5) (Garrido-Martín et al. 2018) with the options ‘--shrink’ and ‘-A median_j’.

### Motif enrichment analysis

The outputs of rMATS for each comparison were used as input to rMAPS (version 2.0.1) (Hwang et al. 2020; Park et al. 2016) in order to identify putative splicing regulators. rMAPS searches for enrichment in binding motifs associated with known RNA binding proteins (RBPs) within the sequences of alternatively spliced transcripts, in particular, in the vicinity of alternatively spliced exons.

### Survivability analysis

Clear cell RCC RNAseq expression data (TCGA-KIRC, 541 samples) and associated metadata were downloaded from the TCGA database (Weinstein et al. 2013) using the ‘TCGAbiolinks’ R package (Colaprico et al. 2016) (version 2.30.0). Kaplan-Meier curves for RBPs, as well as p-values for association between their expression levels and patient survivability, were plotted using the ‘ggsurvplot’ function from the survminer R package (version 0.4.9).

## RESULTS

### Cell deconvolution indicates that kidney tumors lose kidney-specific cell types and gain proliferative characteristics during tumor progression

We downloaded a publicly available bulk RNAseq dataset generated by Wang et al. (X.-M. Wang et al. 2020) from normal kidney samples (N), kidney tumors (C), kidney tumors with thrombi (PT), and their associated thrombi (TT) (Table S1). We then used STAR (Dobin et al. 2013) to perform sequence alignment, and DESeq2 (Love, Huber, and Anders 2014) to obtain the normalized counts matrix (Table S2). We then plotted the samples as data points in PCA latent space (Figure 1A) and observed a clear separation between normal kidney samples (N), tumors without thrombi (C), and thrombi, e.g., thrombi extensions (TT) and their associated tumors (PT). Surprisingly, we could not clearly distinguish between the thrombi extensions (TT) and their associated tumors (PT), indicating that the variability between patients exceeds the differences in gene expression between these two subsets (Figure 1A and Figure S1, see discussion).

**Figure 1:**
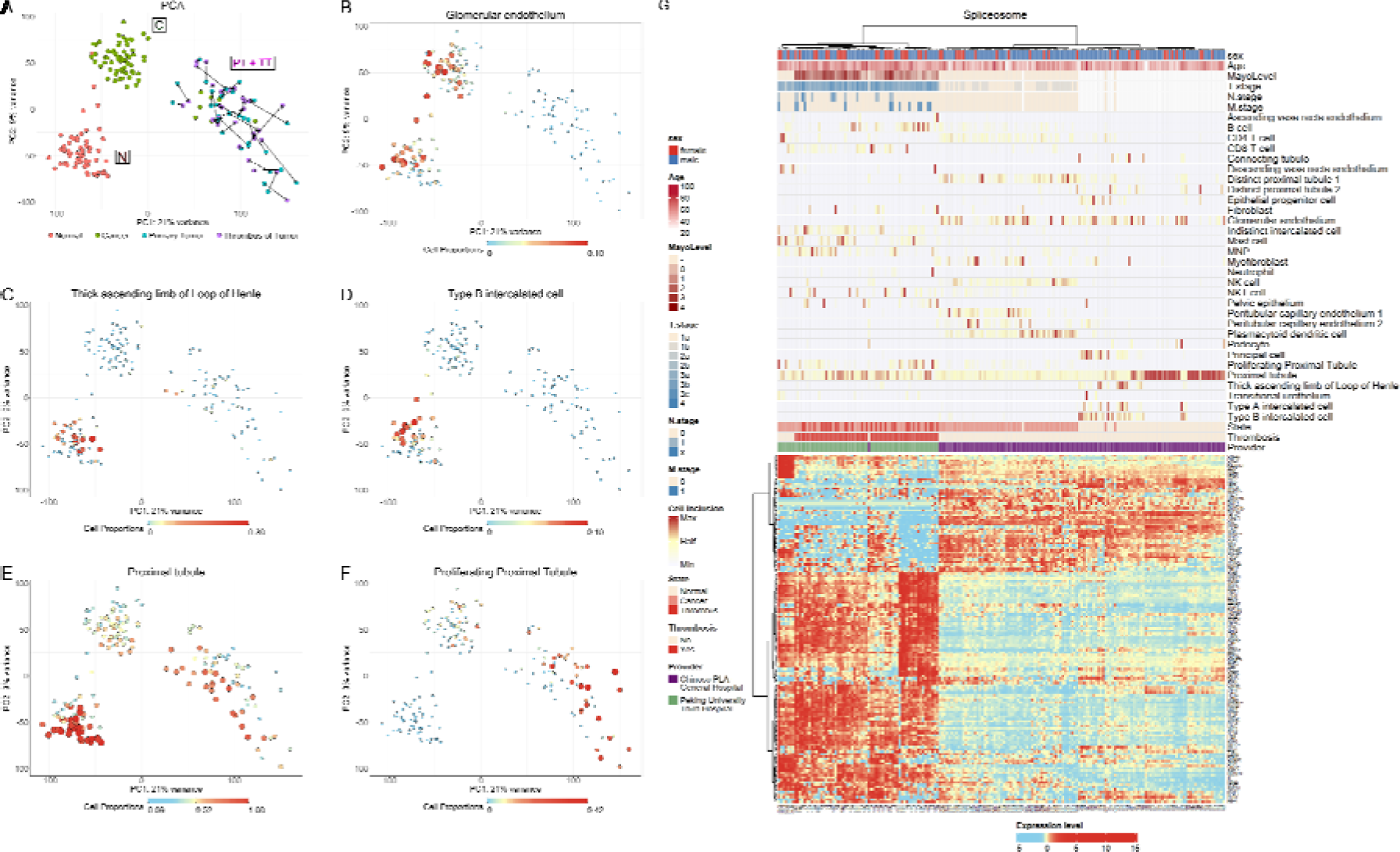
Cell deconvolution indicates that kidney tumors and thrombi are associated with loss of mature kidney-specific cell types and gain of proliferative characteristics. (A) A PCA plot showing gene expression profiles of normal kidney samples (N), kidney tumors (C), kidney tumors with thrombi (PT), and their associated thrombi (TT). The arrows connect between each primary tumor having a thrombus (PT) and its associated thrombus extension (TT). It can be seen that tumors having a thrombus are located close to their associated thrombus in latent space. (B-F) Feature plots showing the proportions of adult kidney cell types within the different samples, as inferred from cell deconvolution. This indicates that although cell proportions of kidney-specific segments (e.g. glomerular endothelium, thick ascending loop of Henle, and type B intercalated cells, and proximal tubule) decrease with tumor formation and thrombus extension, the proportion of proliferating proximal tubular cells is elevated in several tumors with thrombi (PT) and associated thrombi (TT). Dot size and color are proportional to cell proportion (large/red - high, small/blue - low). (G) A clustered heatmap of spliceosome-related genes from HGNC (gene group 1518) along with clinical metadata (top) and cell type proportions from cell deconvolution. It can be seen that the spliceosome related genes distinguish between normal kidney samples, kidney cancer, and kidney tumors with thrombi and their associated thrombi, indicating that splicing is associated with tumor progression and tumor thrombus formation. Note the clear distinction between normal samples presumably originating from the proximal vs. distal segments of the nephron (e.g. LOH and intercalated cells).

We next used BisqueRNA (Jew et al. 2020) to perform cell deconvolution with respect to an adult human kidney single-cell RNA-seq dataset from the Kidney Cell Atlas (Stewart et al. 2019) (Figures 1B-G, Figure S2). Cell deconvolution indicates that most cell proportions of kidney-specific segments (e.g. glomerular endothelium, thick ascending loop of Henle, and type A and B intercalated cells, and the proximal tubule) are lower in the tumors and thrombi relative to normal samples. However, the proportion of proliferating proximal tubular cells were observed to be elevated in a subset of tumors with thrombi (PT) and their associated thrombi (TT) (Figure 1F). Since proliferating proximal tubular cells are also increased following kidney injury (Kusaba et al. 2014; Kusaba and Humphreys 2014) this indicates a distortion in the regeneration and repair mechanisms of the kidney.

### We identified a set of alternatively spliced transcripts that discern between different stages of tumor progression

It is thought that dysregulation of pre-mRNA splicing is a central mechanism in tumor formation and progression (Bonner and Lee 2023; Ivanova et al. 2023; H. Yang, Beutler, and Zhang 2022). In order to understand the relationship between mRNA splicing and thrombus formation in ccRCC, we generated a heatmap of genes related to the major spliceosome (HGNC Gene group: “Major spliceosome”), along with clinical metadata and cell type proportions from cell deconvolution (Figure 1G). We found that the spliceosome-related genes distinguish well between normal kidney samples (N), kidney tumors (C), and the combined set of tumors with thrombi (PT) and their associated thrombi (TT).

In order to find alternatively spliced transcripts related to tumor formation and progression, we used rMATS (Y. Wang et al. 2024) to compare between sets of samples chosen according to their location in latent space (Figure S3). We performed three comparisons: (i) 10 normal samples (N) vs. 10 tumors without thrombi (C); (ii) 10 tumors without thrombi (C) vs. 10 thrombi samples (PT and TT); (iii) 10 normal samples (N) vs. 10 thrombi samples (PT and TT). We also subdivided the thrombi samples into two subsets according to their location in PCA latent space (see Figures S3 and S18). We labeled these two subsets “TH_LOW” and “TH_HIGH” and compared between 12 samples of each group.

We identified a set of transcripts that are alternatively spliced (FDR=0 and |ΔPSI|>0.2, Figure 2, Table S3) between the different normal samples, tumors, and thrombi. For example, the gene ACTN1 has two exons termed 19a and 19b that are alternatively spliced (Figures 2A and S4) (Gardina et al. 2006; Thorsen et al. 2008). It was previously observed that skipping of exon 19b relative to exon 19a is associated with tumor progression (Gardina et al. 2006; Thorsen et al. 2008). In our study we observed that indeed, exon 19b is skipped in thrombi (PT and TT) relative to tumors without thrombi (C) and normal samples (N).

**Figure 2:**
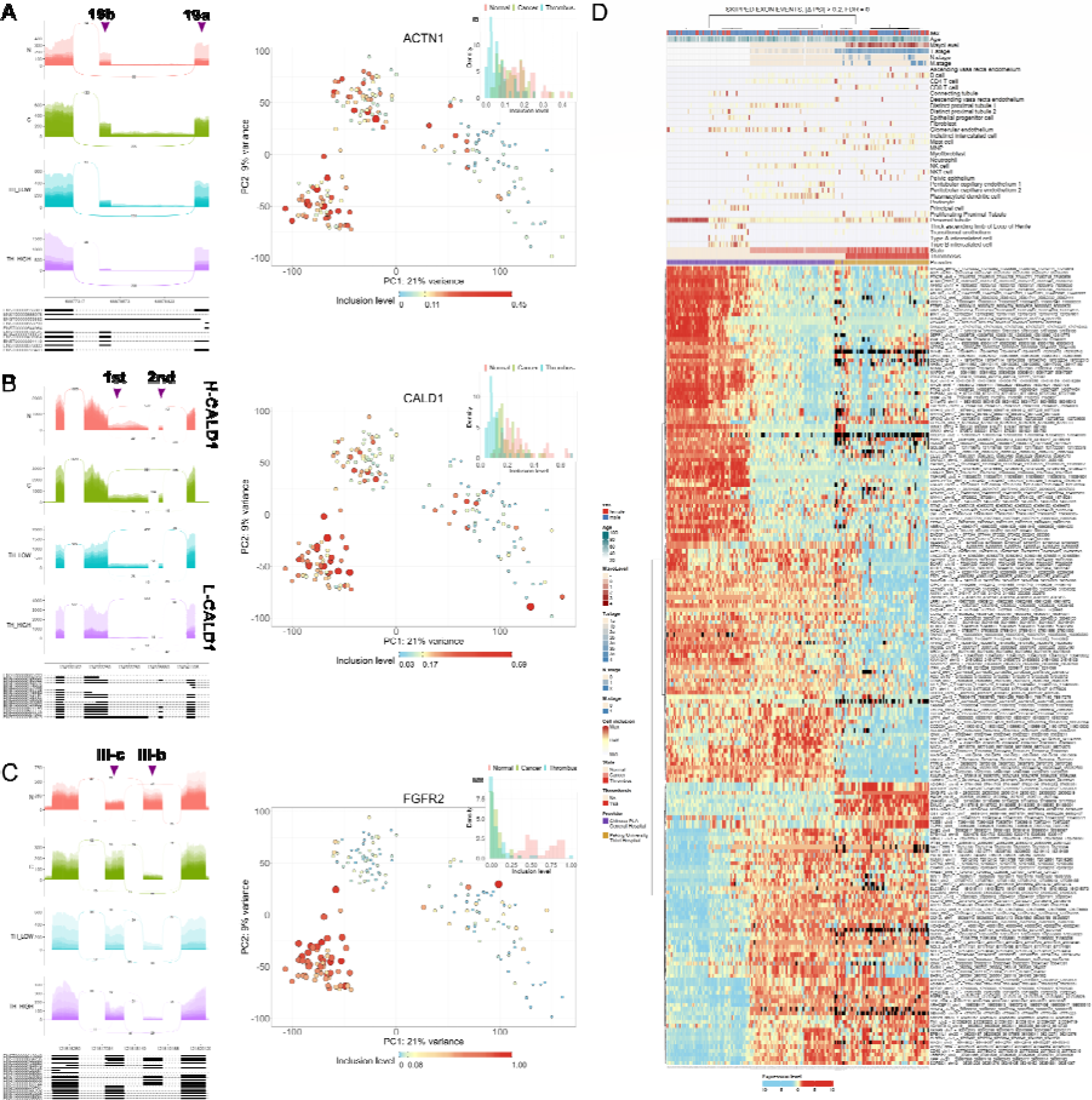
A set of alternatively spliced transcripts discerns between different stages of kidney tumor progression. (A-C) Sashimi and feature plots for selected transcripts that were found to be alternatively spliced between normal kidney samples (N), kidney tumors (C), and thrombi (PT and TT). In the Sashimi plots, the arrow points to the alternatively spliced exon(s). In the feature plots, the color and size of each dot represent the inclusion levels of alternatively spliced exon(s). (D) A heatmap of inclusion levels of transcripts that were found to be significantly alternatively spliced (|ΔPSI| > 0.2, FDR = 0) between normal kidney samples, tumors, and thrombi.

Likewise, the gene CALD1 has two alternatively spliced transcripts (Figures 2B and S5), the heavy isoform H-CALD1, which is the full transcript, and light isoform L-CLAD1, in which two consecutive exons are alternatively spliced such that the 3’ end of the first exon and the full second exon are skipped (Lian et al. 2020; Lin et al. 2009; Thorsen et al. 2008; Yao et al. 2021). It was previously observed that L-CLAD1 is associated with tumor progression and metastasis (Alnuaimi et al. 2022). In our study we observed and over-expression of the L-CALD1 isoform in thrombi (PT and TT) relative to tumors without thrombi (C) and normal samples (N).

The gene FGFR2 has two mutually exclusive exons termed III-b and III-c (Figures 2C and S6) that were previously found to be associated with epithelial and mesenchymal cell phenotypes, respectively (Warzecha, Sato, et al. 2009). Specifically, in kidney tumors it was observed that the epithelial III-b isoform is associated with smaller tumor size, lower tumor grade, and longer patient survival (Zhao et al. 2013). In our study we observed that the epithelial III-b isoform is under-expressed in thrombi (PT and TT) and tumors without thrombi (C) relative to normal samples (N).

The gene HAGH (GLO2) contains 10 exons labeled as “A”, “B”, “1”,…,“8” (Figure 3A and Figure S7). Exon B, that contains a premature termination codon, can be either included or skipped. Transcripts that skip exon B can encode for both mitochondrial and cytosolic isoforms of HAGH, whereas transcripts that include exon B can only encode for cytosolic isoforms (Cordell et al. 2004) (see Figure S7). In our study we observed that exon B has higher inclusion levels in thrombi relative to tumors without thrombi and normal samples, thus effectively encoding mainly for the cytosolic isoform. HAGH (GLO2) is a component of the Glyoxalase system that is responsible for detoxification of methylglyoxal (MGO) and prevents the accumulation of reactive aldehydes in cells (Xue, Rabbani, and Thornalley 2011). It was found that overexpression of the cytosolic Glo2 (cGlo2), but not the mitochondrial Glo2 (mGlo2), inhibited MGO-induced apoptosis of MCF7 breast cancer cells (Xu and Chen 2006). Hence, it is possible that over-expression of the cytosolic isoform (relative to the mitochondrial isoform) in the tumor thrombi may assist in inhibiting cell apoptosis.

**Figure 3:**
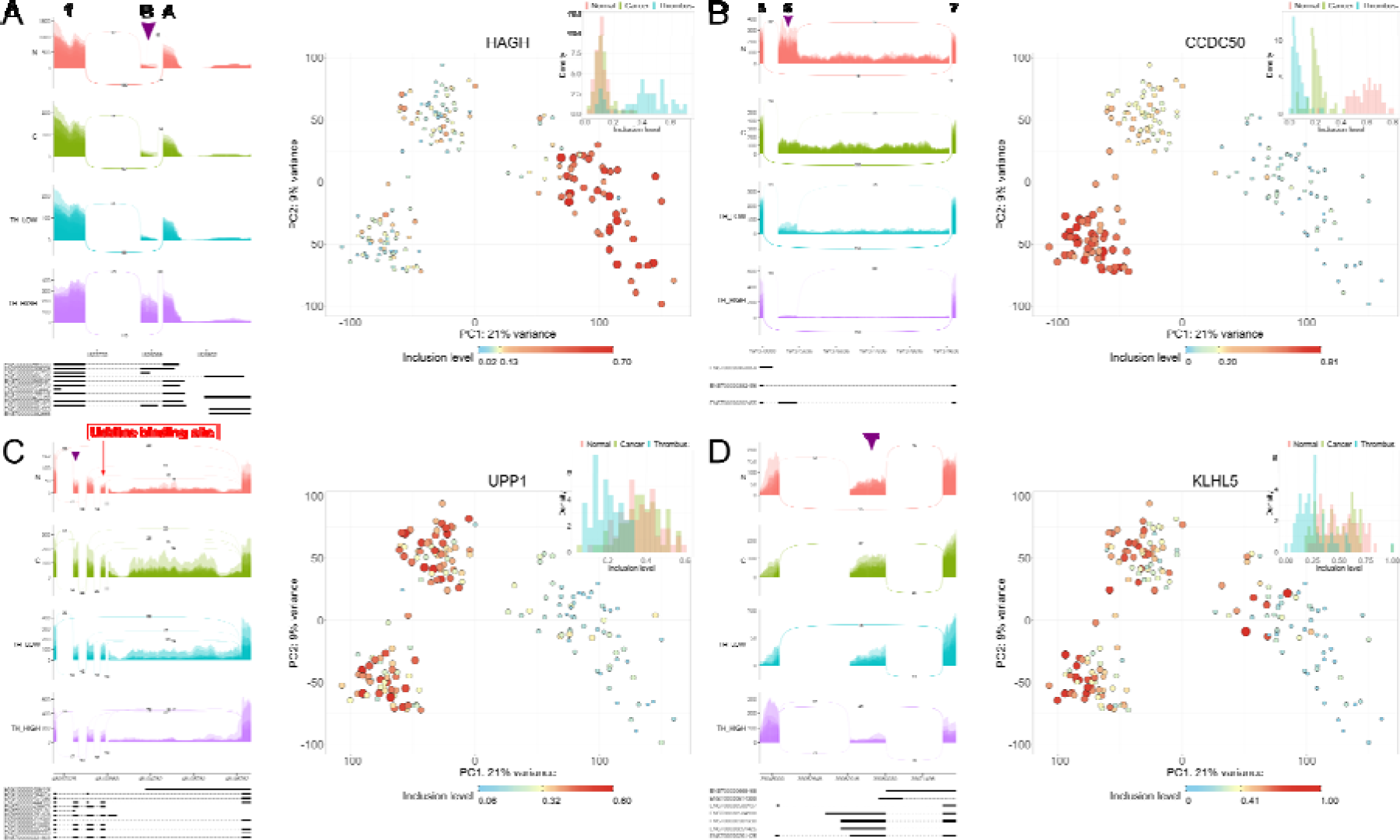
Selected examples of alternatively spliced transcripts that discern between different stages of kidney tumor progression. (A-D) Sashimi and feature plots for selected transcripts that were found to be alternatively spliced between normal kidney samples (N), tumors (C), and thrombi (PT and TT). In the Sashimi plots, the arrows point to the alternatively spliced exon(s). In the feature plots, the color and size of each dot represent inclusion levels of alternatively spliced exon(s).

The gene CCDC50 has two alternatively spliced transcripts (Figure 3B and S9), one long isoform that includes exon 6 (CCDC50L) and one short isoform that skips exon 6 (CCDC50S, NM_174908.3). The short isoform was previously found to be over-expressed in hepatocellular carcinoma and to be correlated with poor tumor differentiation, tumor metastasis, and unfavorable prognosis (H. Wang et al. 2019). In the dataset that we studied we observed that exon 6 of CCDC50 is mostly included in normal kidney samples, moderately skipped in tumors (consistent with Sun et al. (Sun et al. 2020)), and highly excluded in thrombi. We note that the putative splicing regulator SRSF3, that was previously observed to bind and stabilize the CCDC50S isoform (H. Wang et al. 2019), is over-expressed in the thrombi relative to tumors without thrombi and normal samples (Figure S31).

The gene UPP1 (Uridine Phosphorylase 1, UPASE) has a splice variant in which three exons are skipped (NM_001287428.2), thus lacking a uridine binding site (Skinner et al. 2023) (Figures 3C and S8). In the dataset that we studied we observed that UPP1 has higher exclusion of these three exons in thrombi relative to tumors without thrombi and normal samples. We note that it was previously found that disruption of uridine homeostatic regulation in UPP1 knockout mice results in 2–3-fold increase of uridine concentration in the kidney (D. Cao et al. 2005) and leads to spontaneous formation of multiple tumors (Z. Cao et al. 2016).

### Enrichment analysis of known RNA binding motifs identifies putative splicing regulators related to tumor progression

In order to identify splicing regulators, we inserted the results of rMATS obtained earlier from each one of the four comparisons into rMAPS. rMAPS identifies putative splicing regulators by testing for enrichment of RNA binding motifs belonging to known RNA binding proteins (RBPs) in the vicinity of alternatively spliced exons. We found a set of putative splicing regulators that were also differentially expressed between tumors (C) and thrombi (PT and TT), and that showed significant association with survivability in kidney cancer patients from the TCGA KIRC dataset (Weinstein et al. 2013).

We observed that the gene FUS is moderately overexpressed in tumors with respect to normal samples and highly over-expressed in the thrombi (Figure 4A). Independently we observed that high expression of FUS is associated with lower survivability in kidney tumor patients. Given the above, motif enrichment indicates that when highly expressed in thrombi, FUS binds upstream and downstream of target exons and promotes their inclusion. It was previously observed that high expression of FUS in non-small lung cancer is associated with metastasis, large tumor size, and poor prognosis (Xiong et al. 2018). Likewise, it was previously suggested that FUS facilitates the back-splicing of circEZH2, a transcript which promotes tumorigenesis, EMT, and metastasis in breast cancer (Liu et al. 2022).

**Figure 4:**
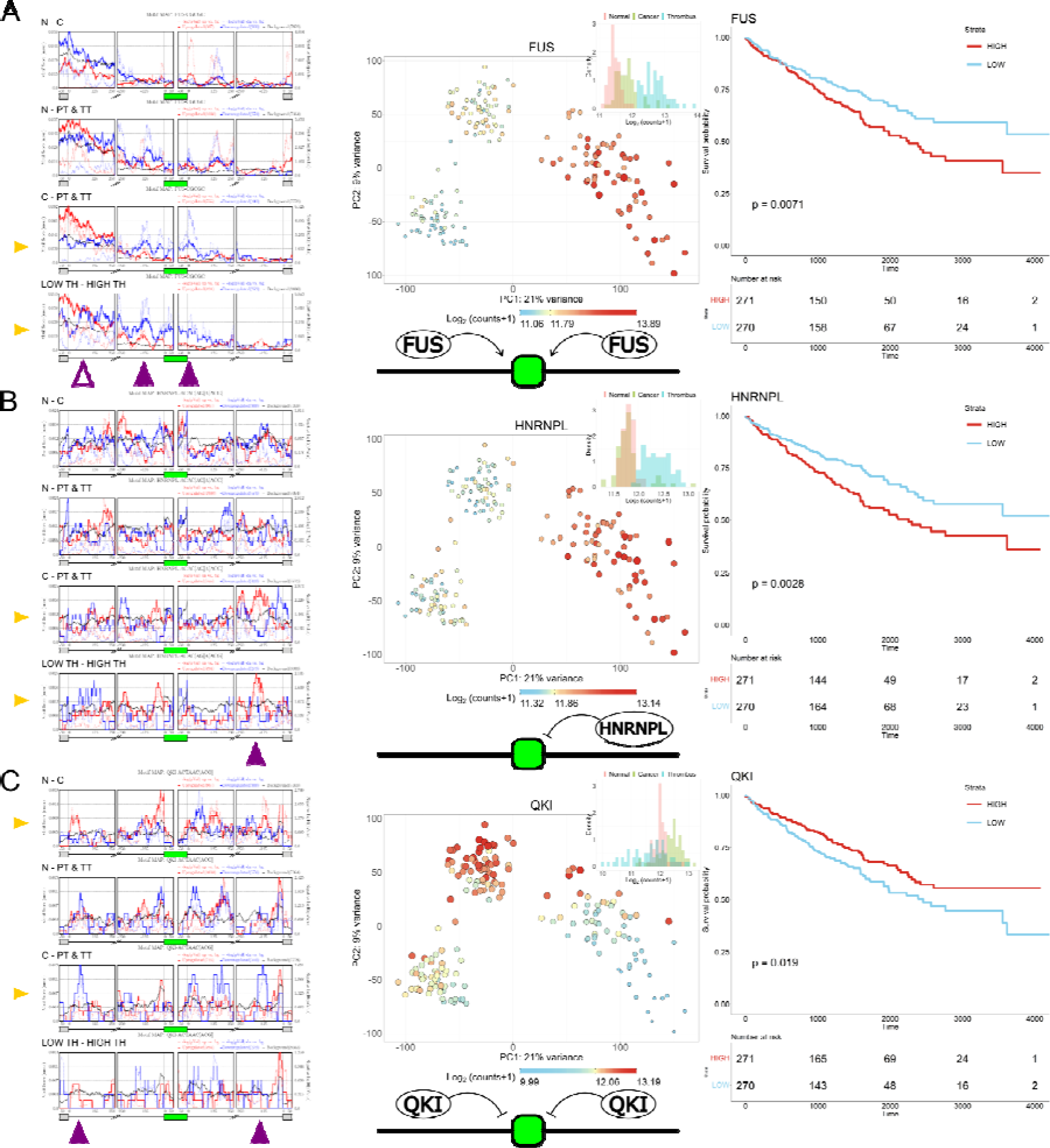
Enrichment analysis for known RNA binding motifs identifies putative splicing regulators related to kidney tumor progression. (A-C) Left panels shows motif enrichment plots for selected RNA binding proteins (RBPs). Middle panels show feature plots for RBP expression levels (red/large - high expression, blue/small – low expression). Right panels show Kaplan-Meier curves obtained for each selected RBP from the from TCGA clear-cell RCC dataset (KIRC). It can be seen that over-expression of the RBPs FUS and HNRNPL is associated with tumor thrombi and lower survivability. Presumably, in the tumors with thrombi, FUS binds upstream and downstream of cassette exons and promotes their inclusion, whereas HNRNPL binds downstream and promotes their exclusion. On the other hand, QKI which is associated with slightly higher survivability, is over-expressed in the tumors without thrombi and presumably binds upstream and downstream of cassette exons and promotes their exclusion.

Likewise, we observed that the gene HNRNPL has a higher expression in the thrombi with respect to tumors and normal samples, and that high expression of HNRNPL is associated with lower survivability in kidney tumor patients (Figure 4B). Together with motif enrichment, this indicates that when highly expressed in the thrombi, HNRNPL binds downstream of target exons and inhibits their inclusion. It was previously observed that HNRNPL is highly expressed in liver cancer, pancreatic cancer, prostate cancer, and a variety of tumors, and promotes tumor proliferation, migration and invasion (Gu et al. 2020). It was previously observed in prostate cancer that HNRNPL forms a complex with SNHG1, a long non-coding RNA which promotes EMT and metastasis (Tan et al. 2024). SNHG1 is also upregulated in the ccRCC thrombi (Figure S30).

We also observed that the gene QKI has a lower expression in normal samples with respect to cancer, and even lower expression in a subset of thrombi (Figure 4C). Independently, we observed that low expression of QKI is associated with lower survivability in kidney tumor patients. Taken together with motif enrichment analysis, this indicates that when highly expressed in the cancer samples (C, without thrombi), QKI binds both upstream and downstream of target exons and inhibits their inclusion. We note that it was previously observed that QKI can promote mesenchymal splicing patterns and increased tumorigenicity (Li et al. 2018; Y. Yang et al. 2016).

## DISCUSSION

In this study we set to characterize alternative splicing in ccRCC thrombi using a publicly available RNAseq dataset. We first projected the samples to latent space and used cell deconvolution to show that a subset of thrombi and their associated tumors had higher proportions of cells resembling proliferating proximal tubular cells. Likewise, we observed that expression levels of spliceosome-related genes discern between normal samples, tumors, and thrombi. We then found a set of transcripts that are alternatively spliced between normal samples, tumors, and thrombi, and also identified putative splicing regulators whose expression is associated with patient survivability. These findings are consistent with previous indications that dysregulated mRNA splicing, such as abnormal expression of spliceosome genes or mutations in components of the splicing machinery, are associated with tumor formation and progression (Deng et al. 2021; Ivanova et al. 2023; Niño, Scotto di Perrotolo, and Polo 2022; Zhang et al. 2021). We believe that our research can lead to better understanding of mRNA splicing mechanisms underlying thrombus formation in ccRCC, and eventually assist in the development of RNA-based therapeutics (Havens and Hastings 2016; Ishigami et al. 2024; J. Kim et al. 2023; Tang et al. 2021).

We observed that the tumors that have thrombi (PT) cluster together with their associated thrombi (TT) in PCA latent space and not with tumors lacking thrombi (C) (Figure 1A). One possible explanation for this is that tumors having thrombi (PT) and their associated thrombi (TT) have significantly different genomic characteristics from tumors lacking thrombi (C) due to early mutation events associated with thrombus formation (X.-M. Wang et al. 2020). As a cautionary note, it is important to keep in mind that different medical centers provided the tumors with thrombi (PT and TT) and the tumors lacking thrombi (C) (Figure 1G).

Although there exist single-cell RNAseq datasets from ccRCC thrombi (Shi et al. 2022), it is difficult to use them to characterize heterogeneity between patients since only a relatively small number of samples are available due to the complexity and cost of these experiments. Moreover, since they are based on short-read technologies, whereby a cell barcode and UMI are inserted during 3’ end priming, they do not cover full transcript lengths, and are therefore less suitable for identifying alternative mRNA splicing. Recent developments in long-read sequencing technologies (Al’Khafaji et al. 2024) will likely enable deeper investigation of alternative mRNA splicing at the single-cell level.

## Supporting information

Table S1 - metadata

Table S2 - raw and normalized counts

Table S3 - inclusion levels 3 comparisons

## ACKNOWLEDGEMENTS

We wish to thank Achia Urbach and all members of our lab for helpful comments and suggestions.

## AUTHOR CONTRIBUTIONS

Study initiation and conception – Z.C. and T.K.; Data analysis - Z.C. and T.K.; Other intellectual contribution – H.R. and I.B.D.; Manuscript writing – Z.C. and T.K.; All authors have read and agreed to the published version of the manuscript.

## DATA AVAILABILITY STATEMENT

The results shown here are in part based upon data generated by the TCGA Research Network: https://www.cancer.gov/tcga.

## APPENDICES

**Supplementary information:** Supplementary text and figures.

**Table S1:** A table of sample metadata.

**Table S2:** A table of raw and normalized gene expression counts.

**Table S3:** A table of inclusion levels of genes that were found to be significantly alternatively spliced (skipped exons with |ΔPSI| > 0.2 and FDR = 0) between normal kidney samples (N), tumors (C), and thrombi (PT and TT).

**Program:** A compressed directory containing programs and datasets for data visualization.

