## Supplementary text and figures. for "Characterization of alternative mRNA splicing associated with tumor thrombus in clear-cell renal cell carcinoma"

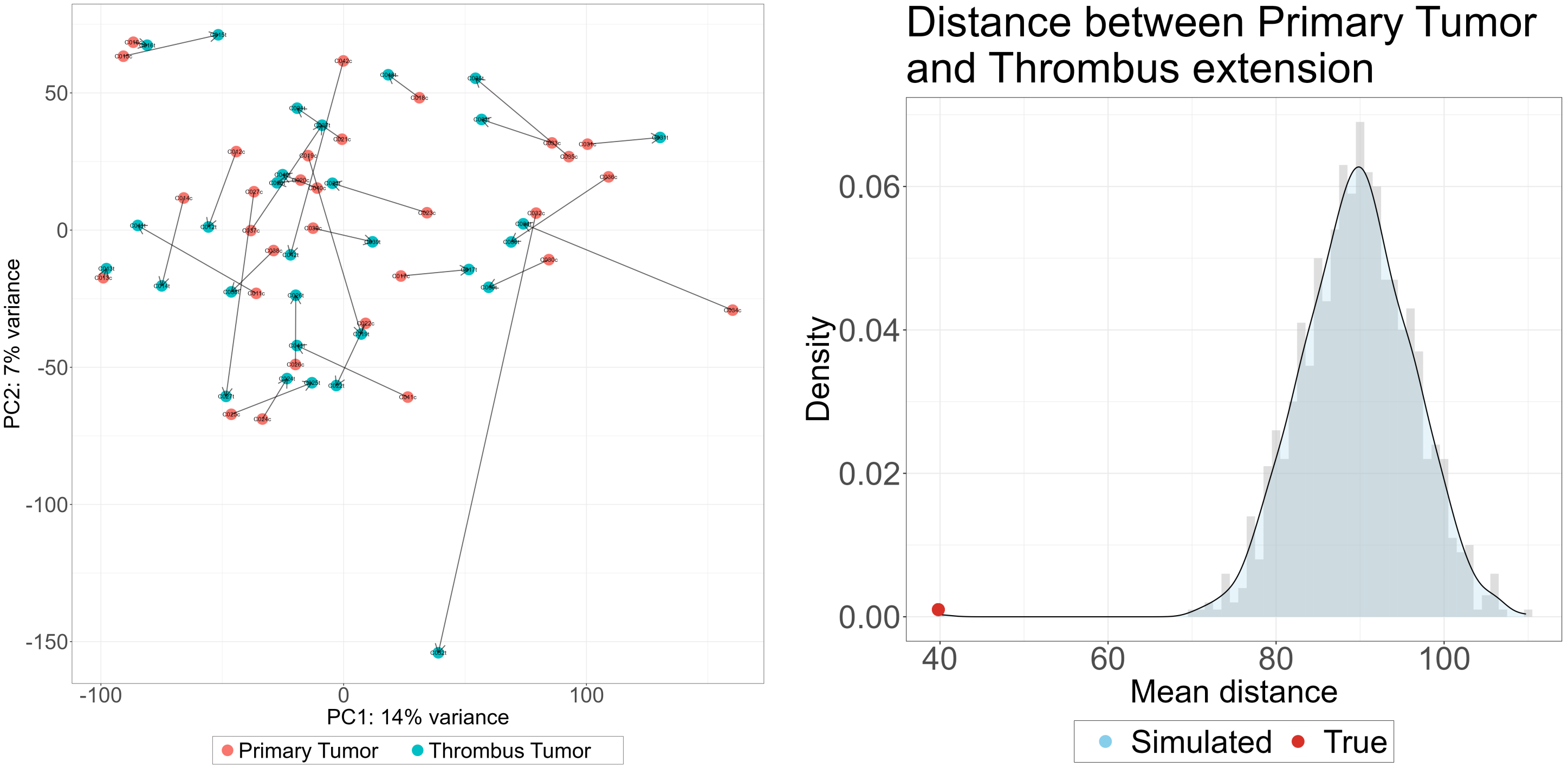

**B**

**A**

### Figure S1: The heterogeneity between patients is larger than the difference between tumors (PT) and their associated thrombi (TT)

(A) A PCA plot of 30 primary tumors and associated thrombi, where each tumor and its associated thrombus (i.e., paired samples from the same patient) are connected with an arrow. (B) The mean distance between each tumor (PT) and its associated thrombus (TT) in latent 2D PCA space (which we define as the mean PT-TT distance) is much smaller than the distance between tumors and thrombi in randomized pairing configurations. Shown is the distribution of the mean PT-TT distance in 999 configurations, where each configuration consists of random pairing between tumors and thrombi. It can be seen that the actual mean PT-TT distance is significantly lower (Z = -7.38) than the randomized mean PT-TT distances.

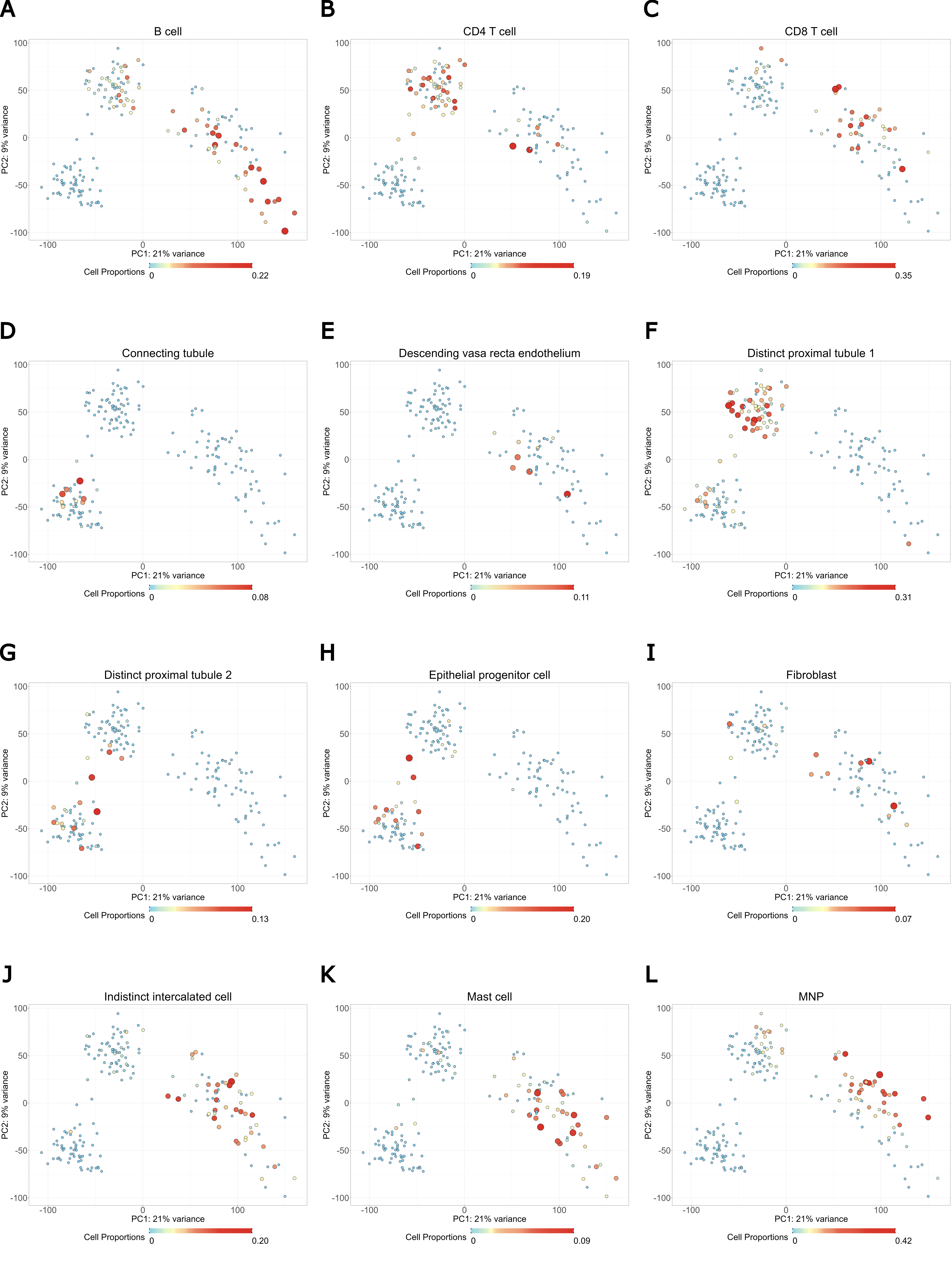

Continues in next page.

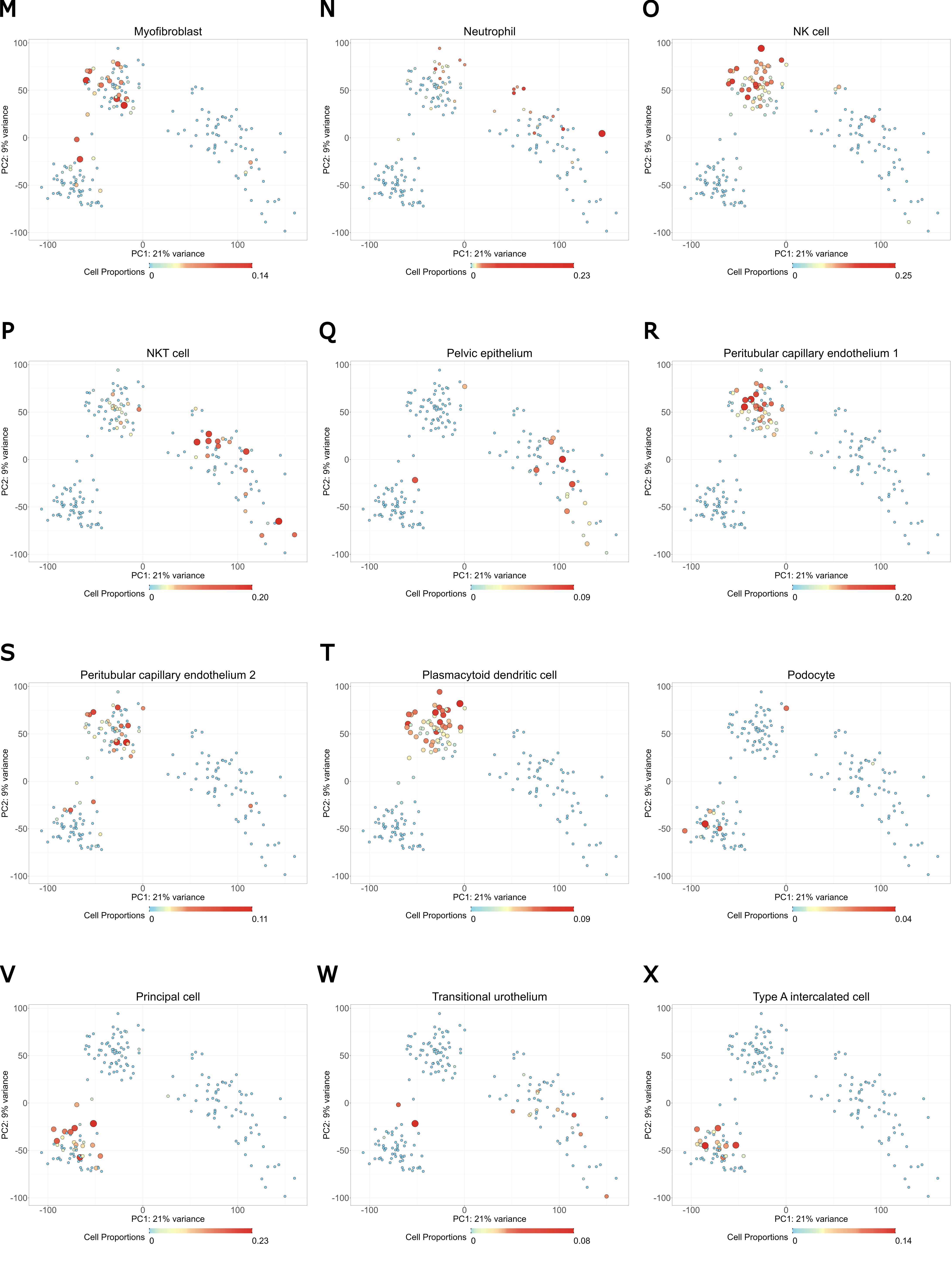

U

### Figure S2: Feature plots of cell type proportions across the different samples.

(A-X) Cell type proportions estimations using BisqueRNA deconvolution. Dot size and color are proportional to cell proportion (large/red - high, small/blue - low). We omitted podocytes due to low proportions across most of the samples. All proportions are also shown in the heatmap in Figure 1G. MNP – mononuclear phagocytes.

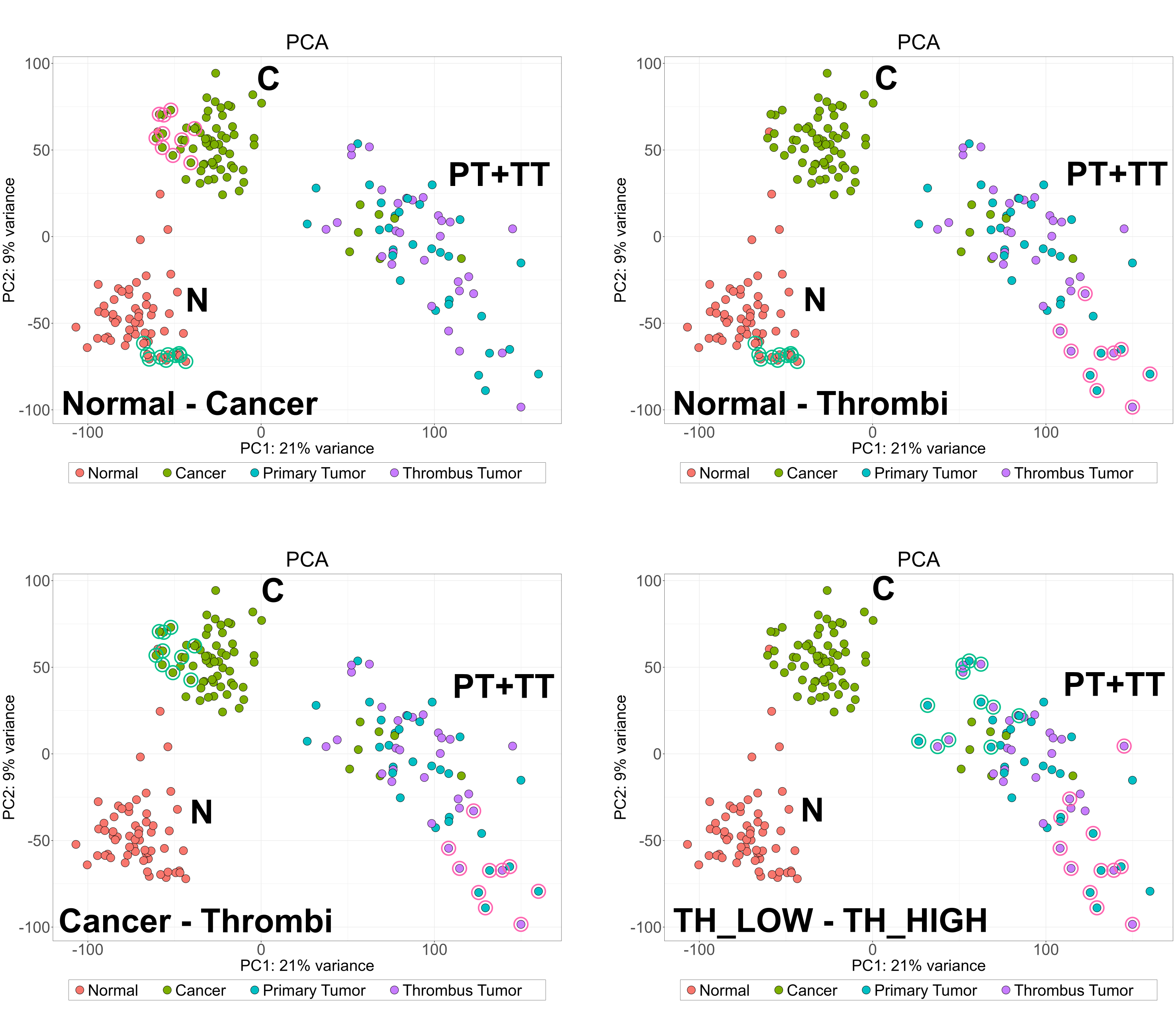

A

C

B

D

### Figure S3: PCA plots marking samples that were used for rMATS comparisons.

rMATS was used to compare representative samples from each group. Shown are PCA plots in which the samples used for the rMATS comparisons are circled. Here the first representative group is encircled in green, and the second representative group is encircled in red. Representative samples were chosen based on position in PCA latent space. See also Figure S18.

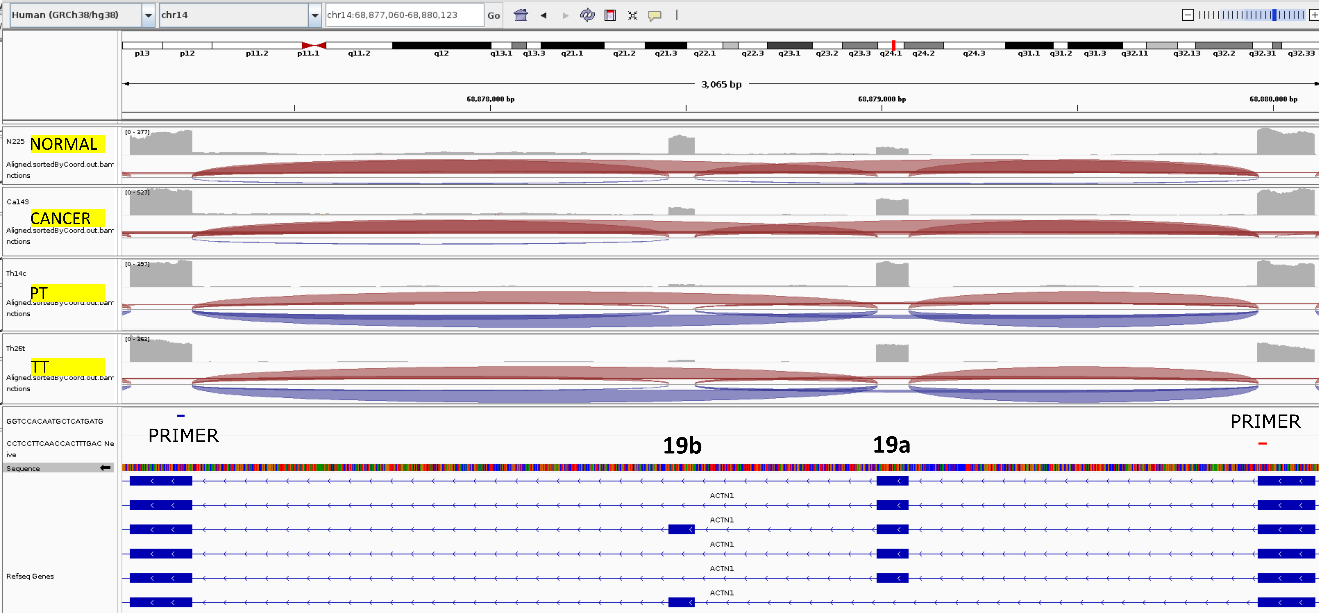

### Figure S4: Alternative splicing in the gene ACTN1.

Locations of exons were verified based on primers from (Gardina et al. 2006).
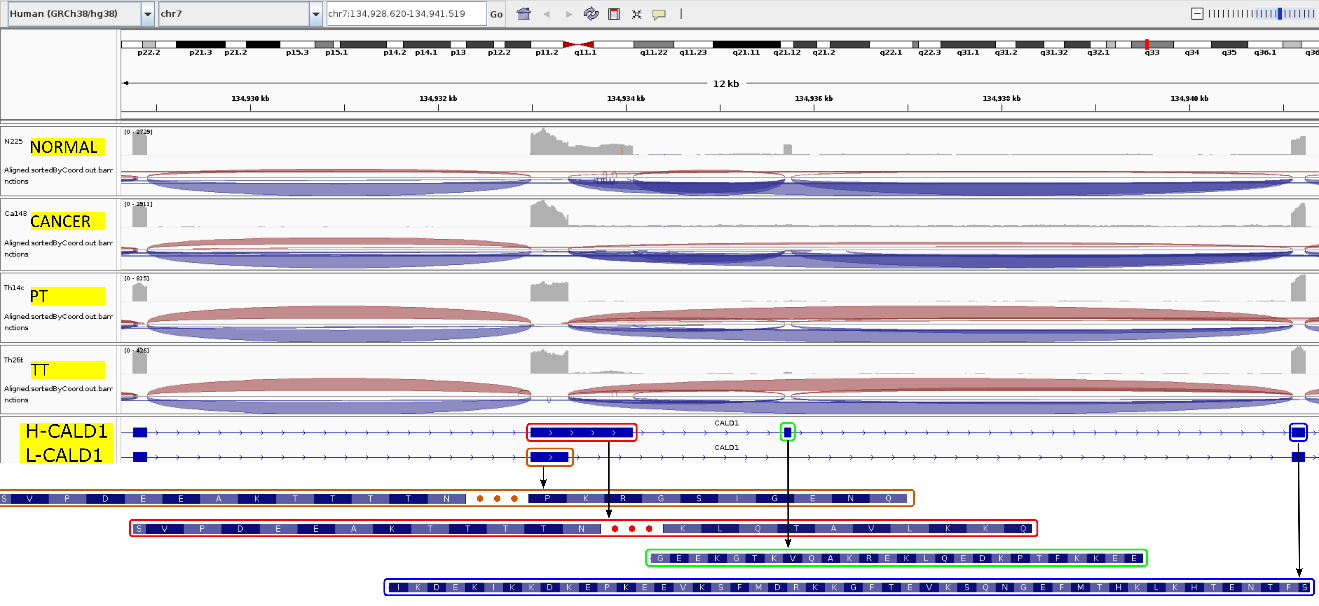

### Figure S5: Alternative splicing in the gene CALD1.

Identity of exons was verified based on amino acid sequences found in (Lin et al. 2009, 1).

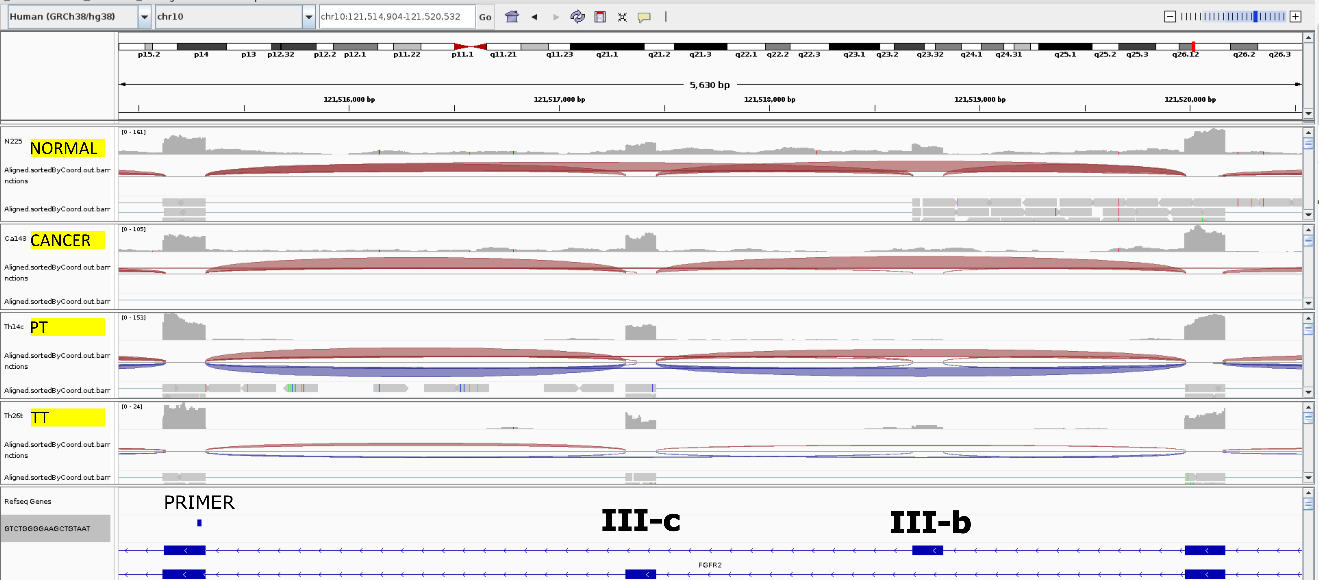

### Figure S6: Alternative splicing in the gene FGFR2.

Locations of exons were verified based on a primer from (Scotet and Houssaint 1998).

A

B

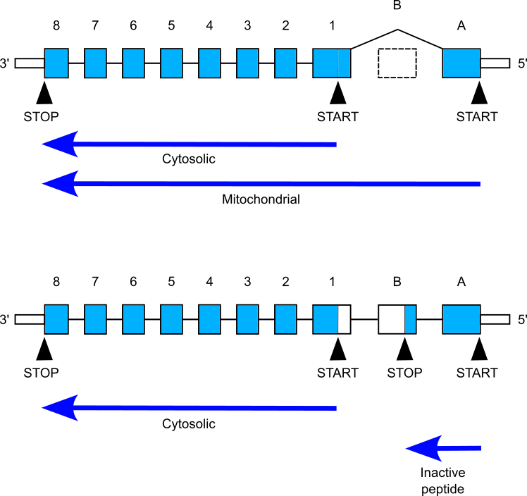

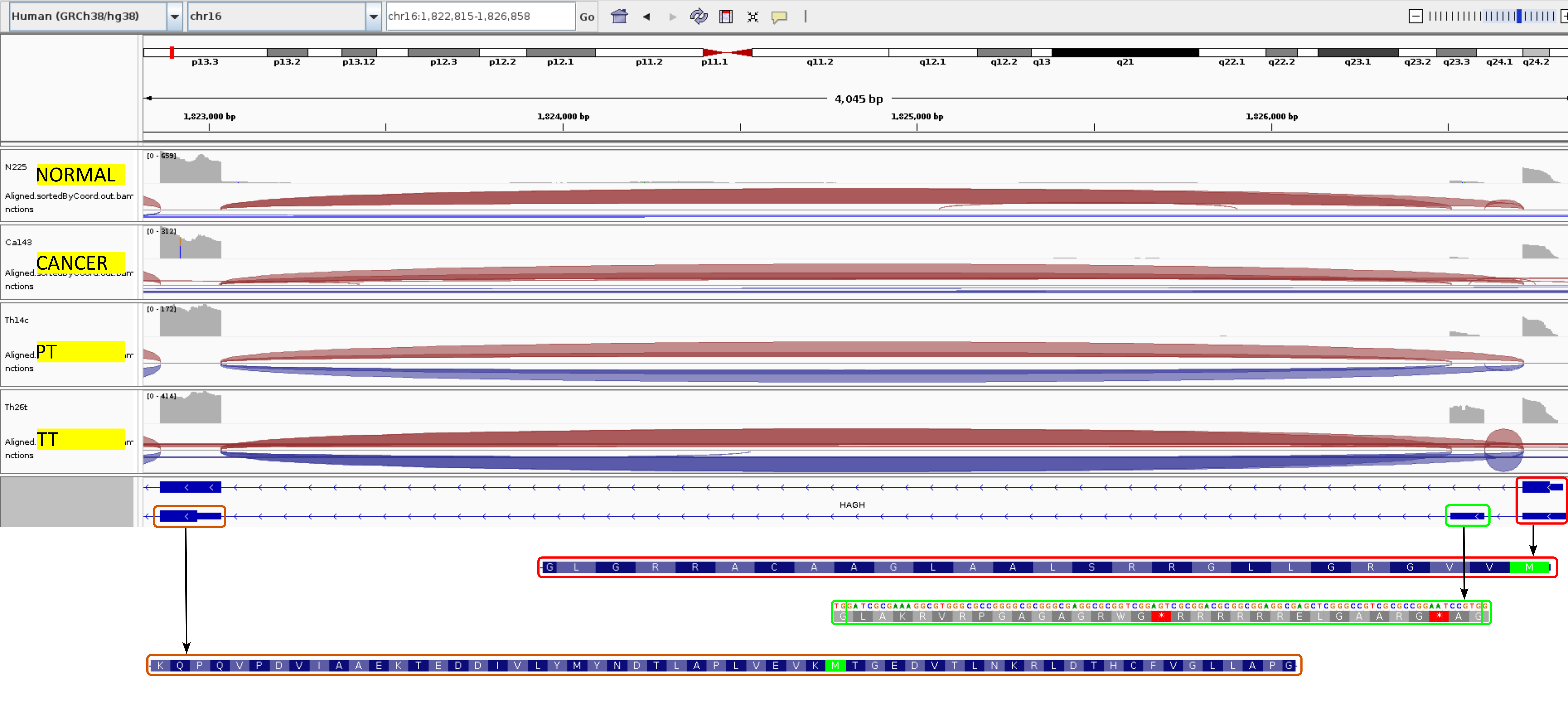

**A**

**B**

**1**

### Figure S7: Alternative splicing in gene HAGH.

Identity of exons was verified based on amino acid sequences found in (Cordell et al. 2004).

A

B

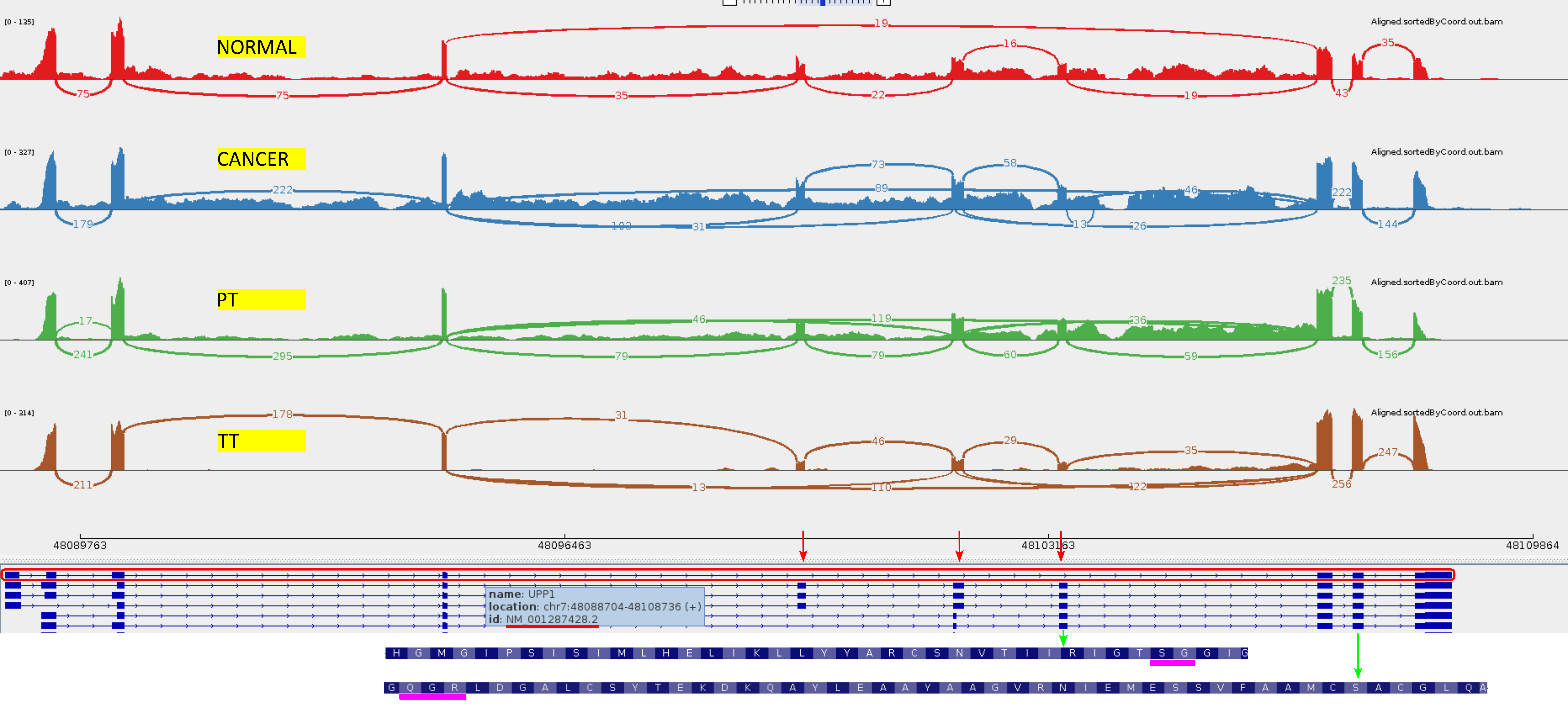

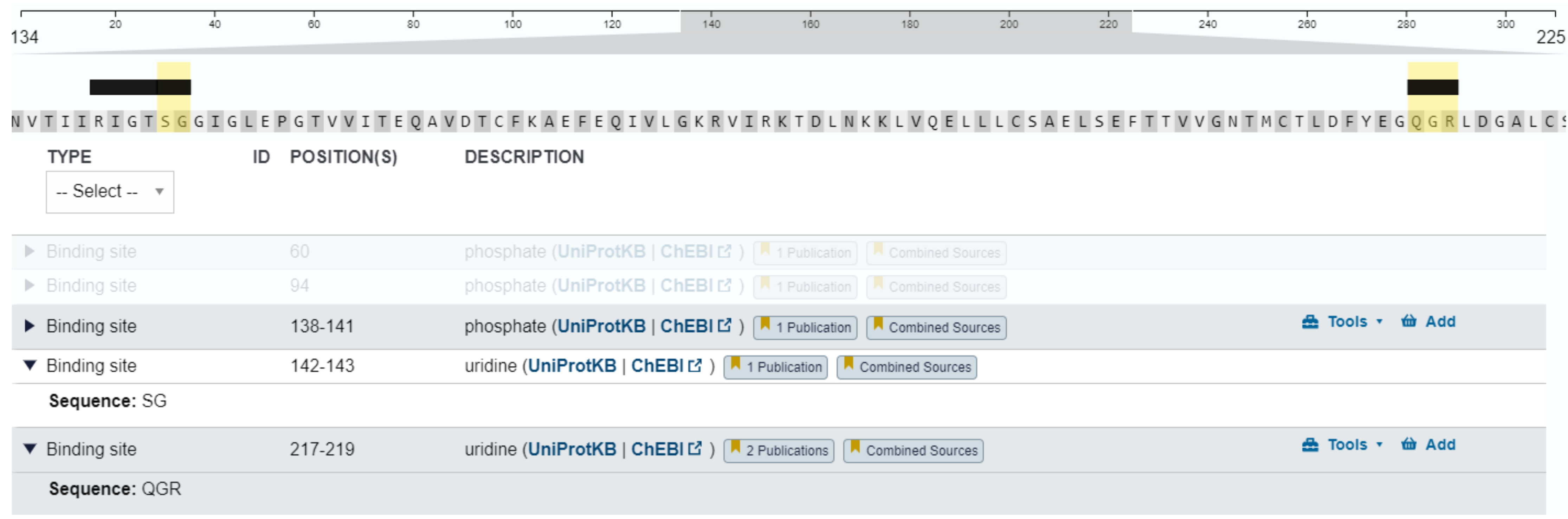

### Figure S8: Alternative splicing in the gene UPP1.

Locations of exons were verified based on the transcript ID (NM_001287428, marked in red) found in (Skinner et al. 2023). Uridine binding sites were identified with UniProt (The UniProt Consortium 2023).

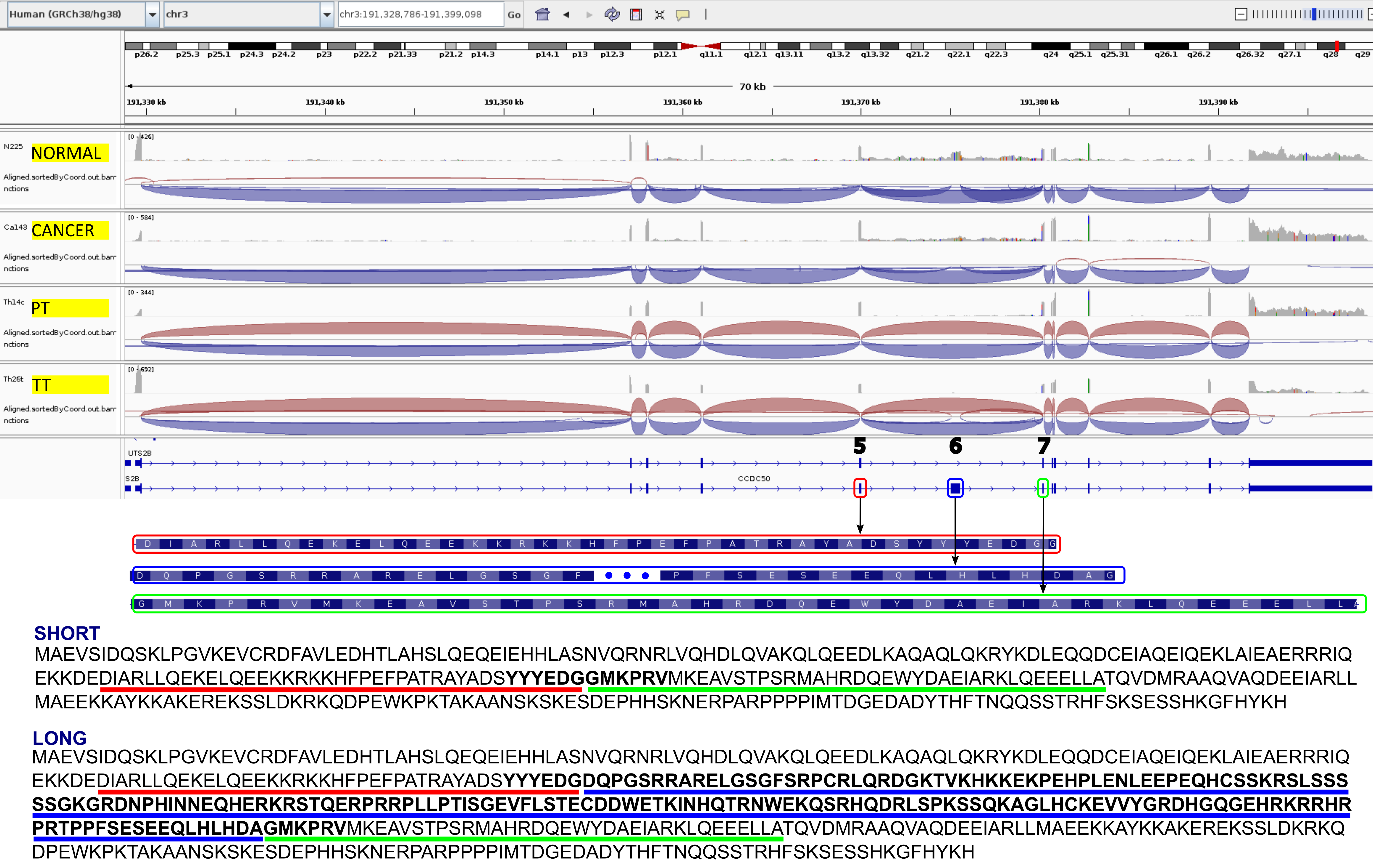

**A**

**B**

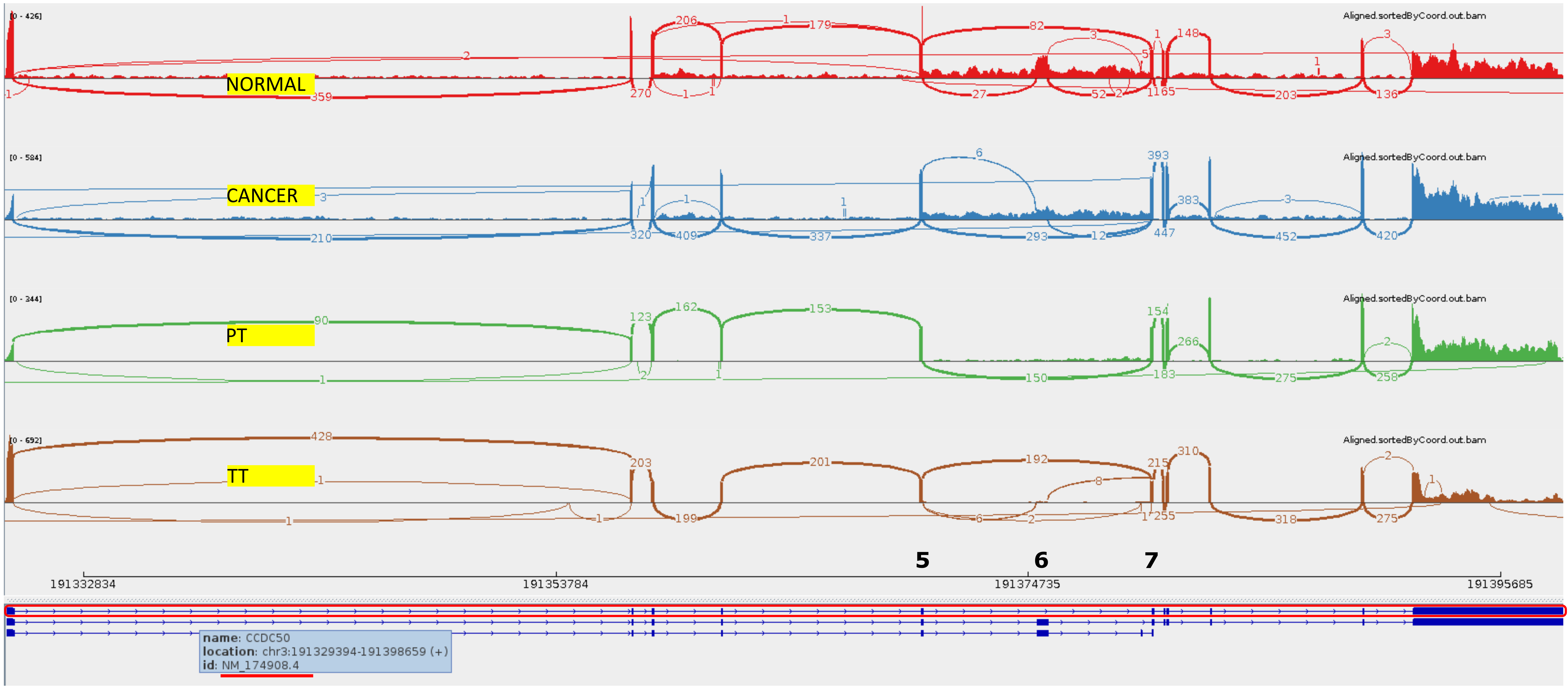

### Figure S9: Alternative splicing in the gene CCDC50.

Locations of exons were verified based on transcript ID found in (Wang et al. 2019) and amino acid sequence found in (Vazza et al. 2003).

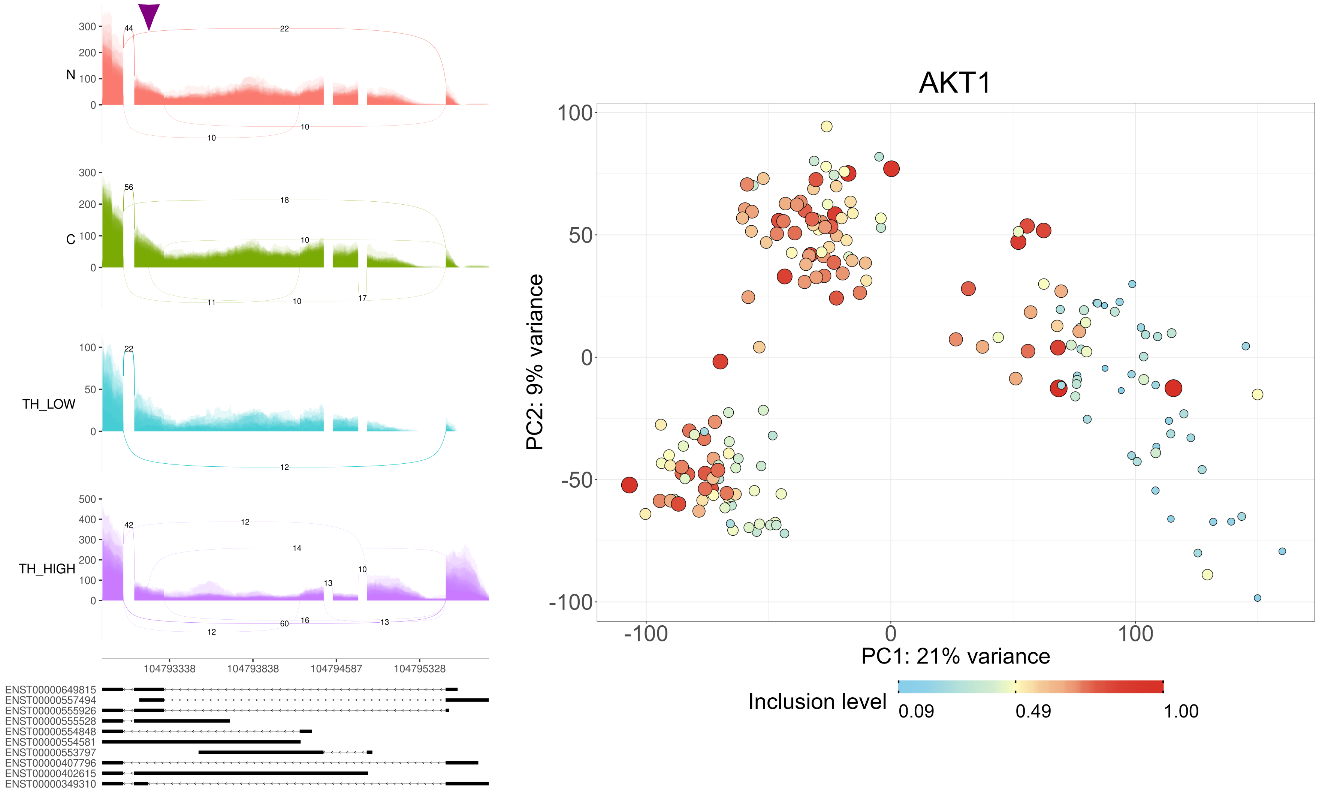

### Figure S10: Alternative splicing in the gene AKT1.

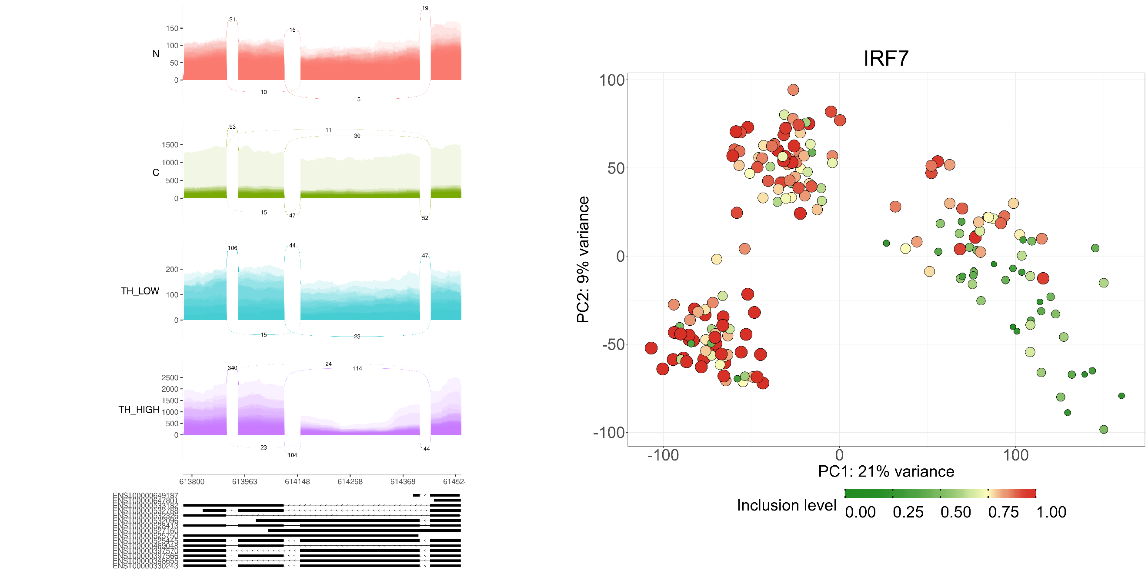

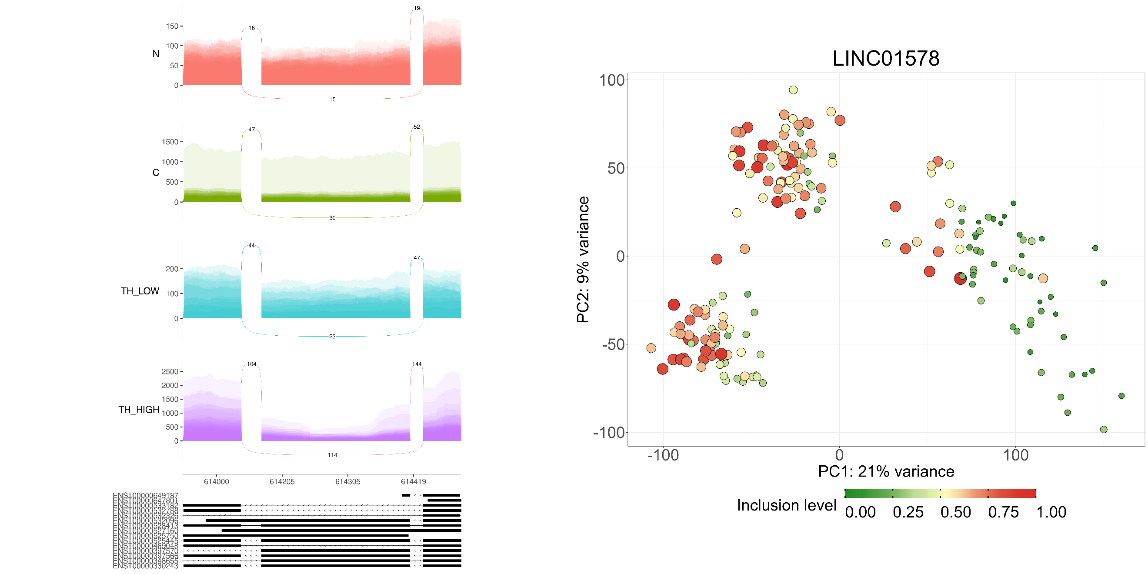
Sashimi and feature plots of inclusion levels for a cassette exon in the gene AKT1.

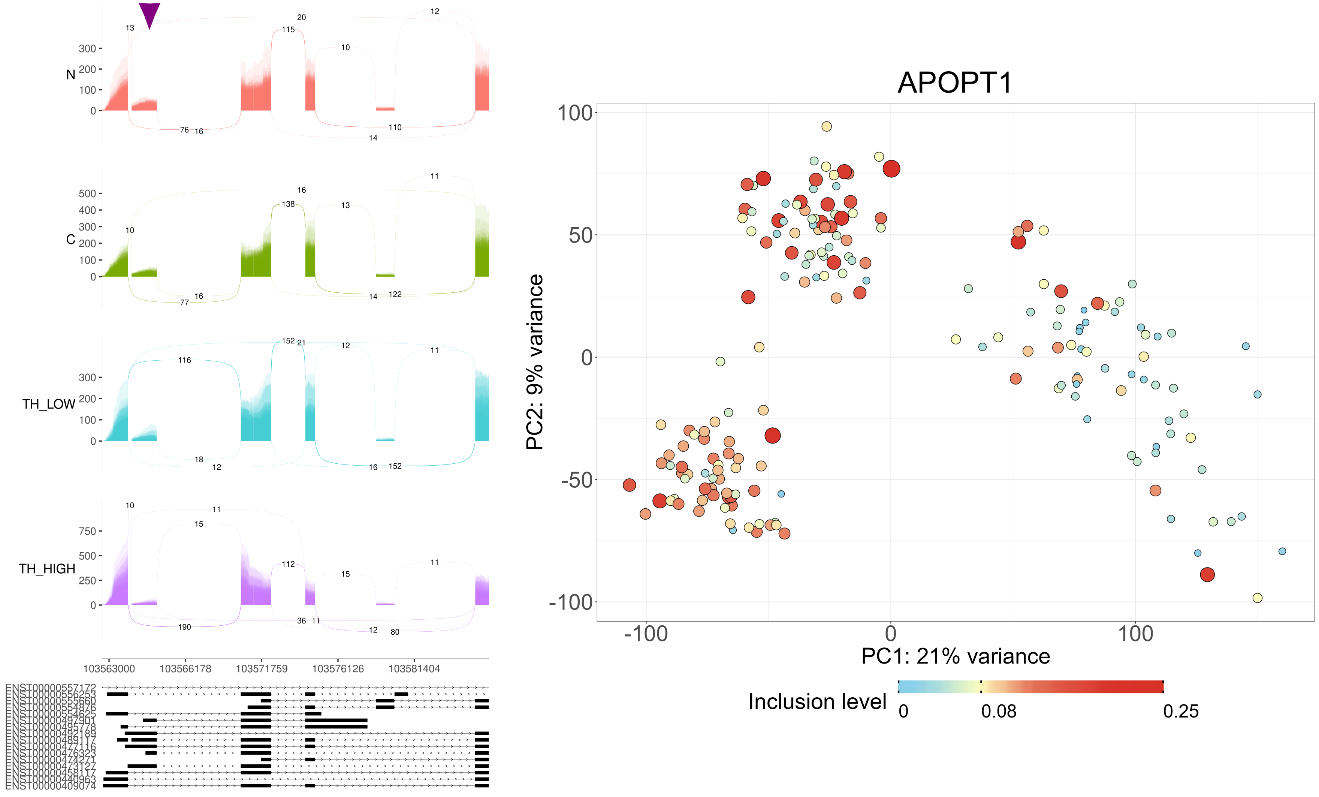

### Figure S11: Alternative splicing in the gene APOPT1.

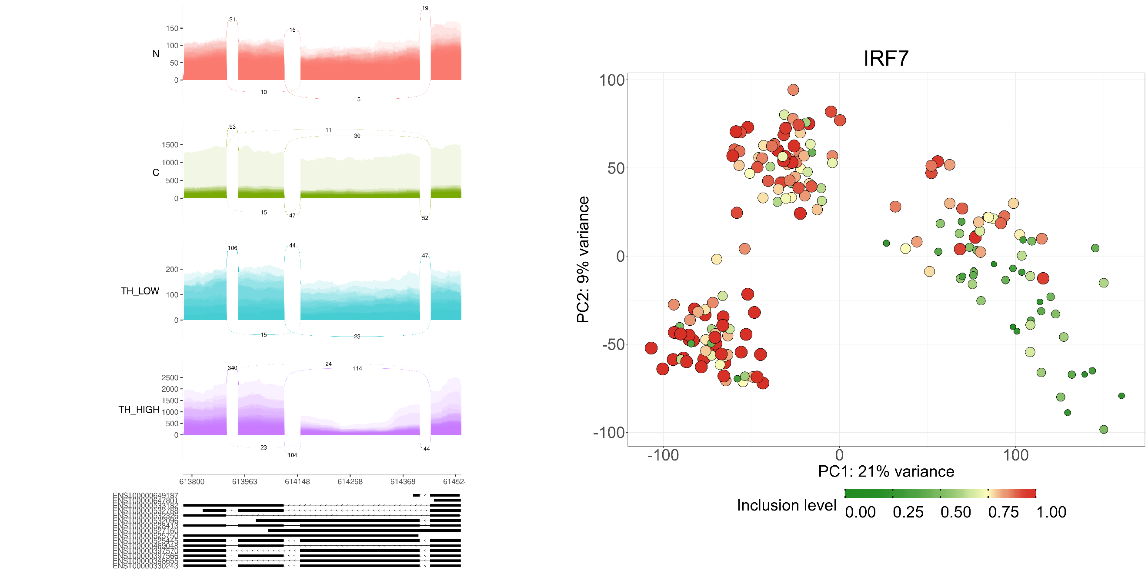

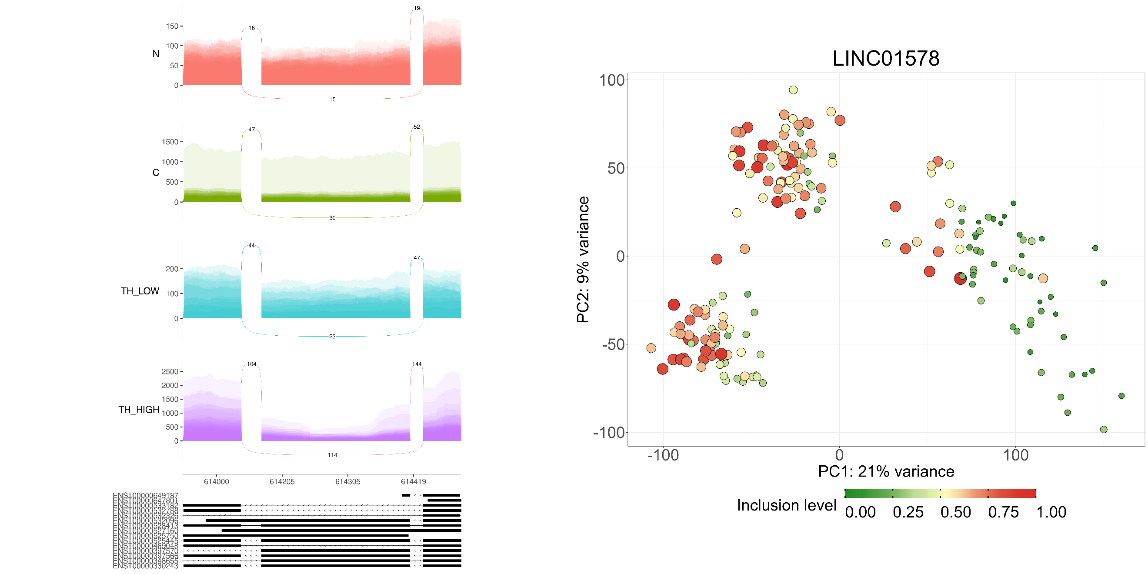
 Sashimi and feature plots of inclusion levels for a cassette exon in the gene APOPT1.

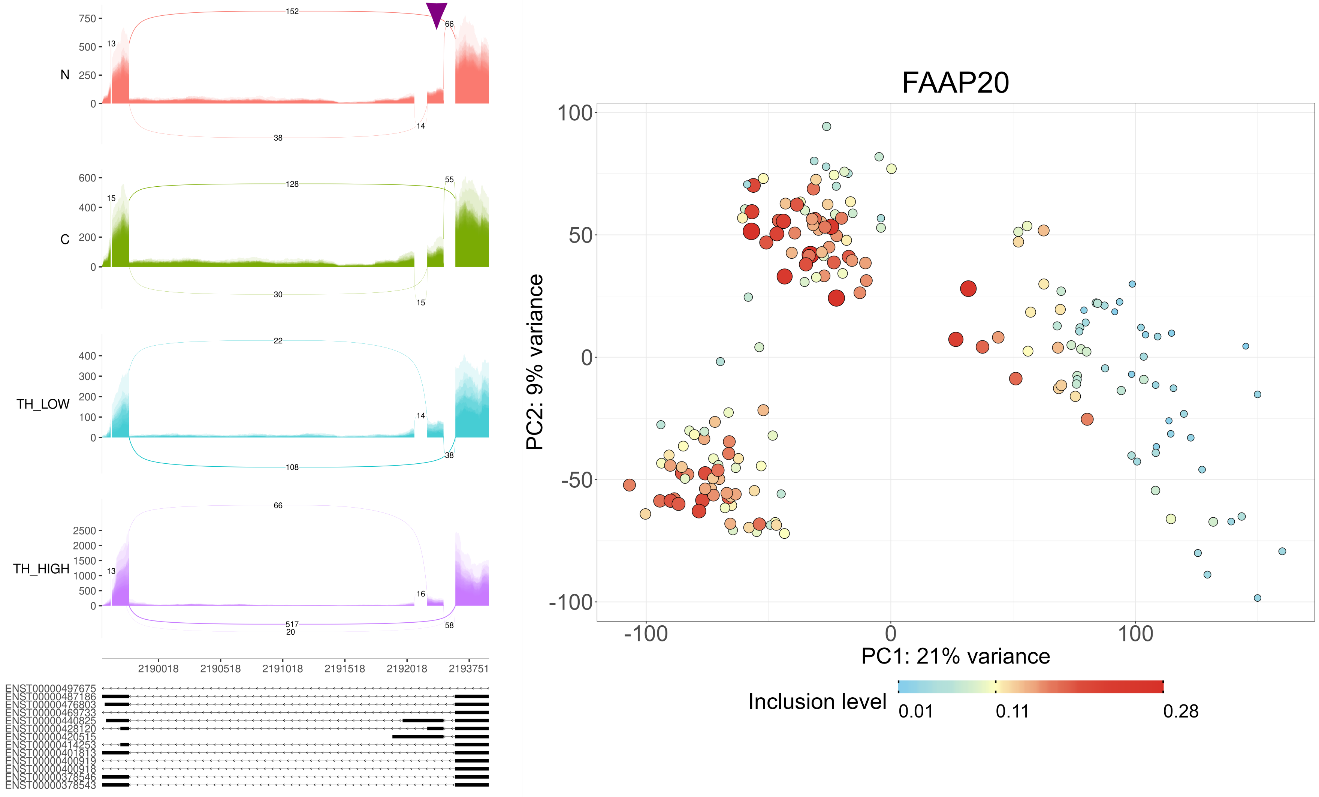

### Figure S12: Alternative splicing in the gene FAAP20.

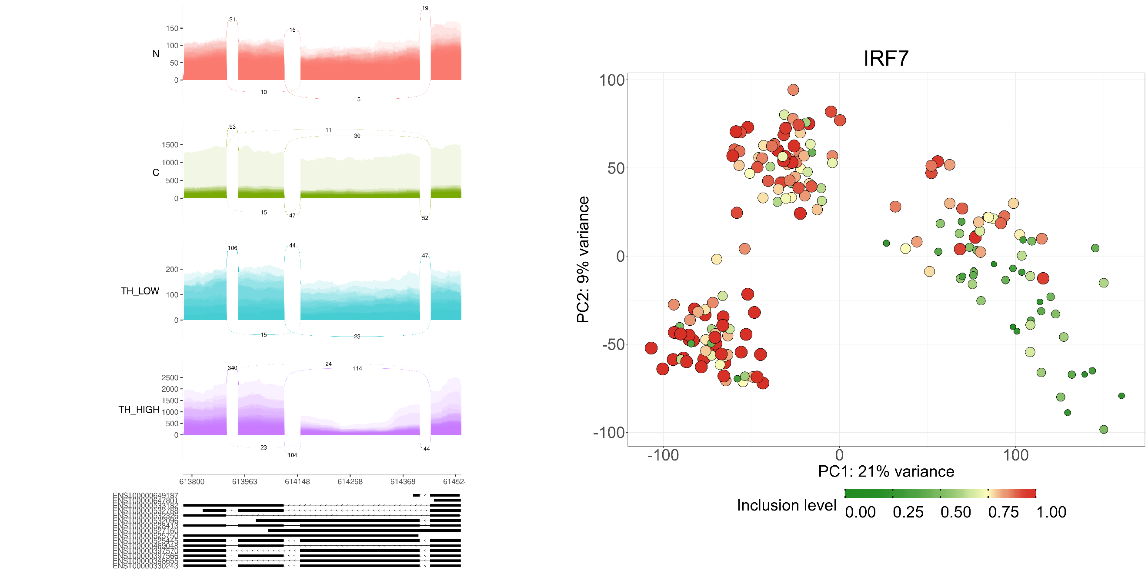

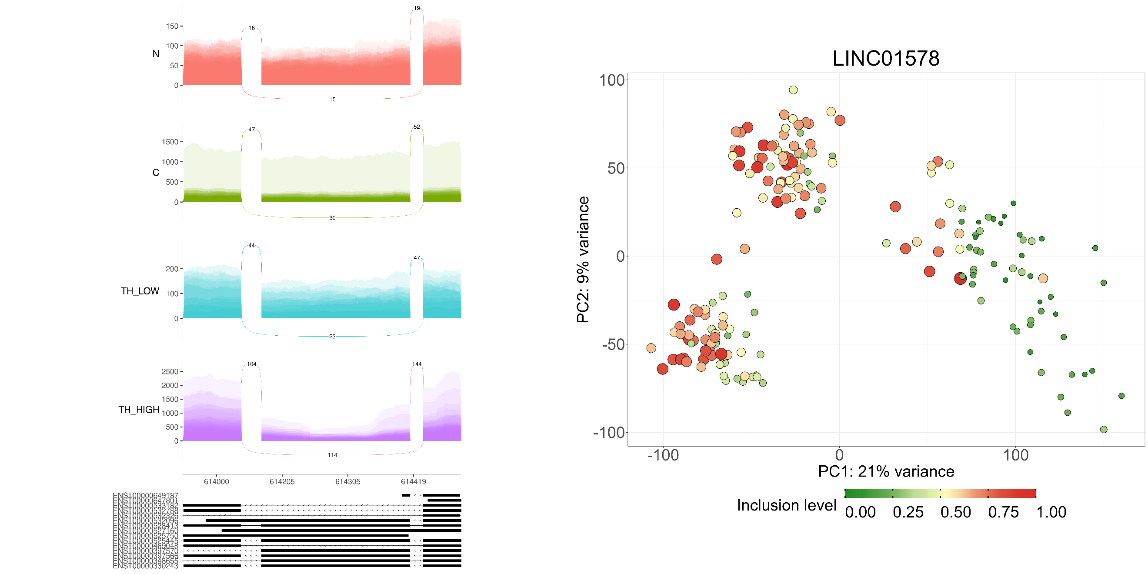
 Sashimi and feature plots of inclusion levels for a cassette exon in the gene FAAP20.

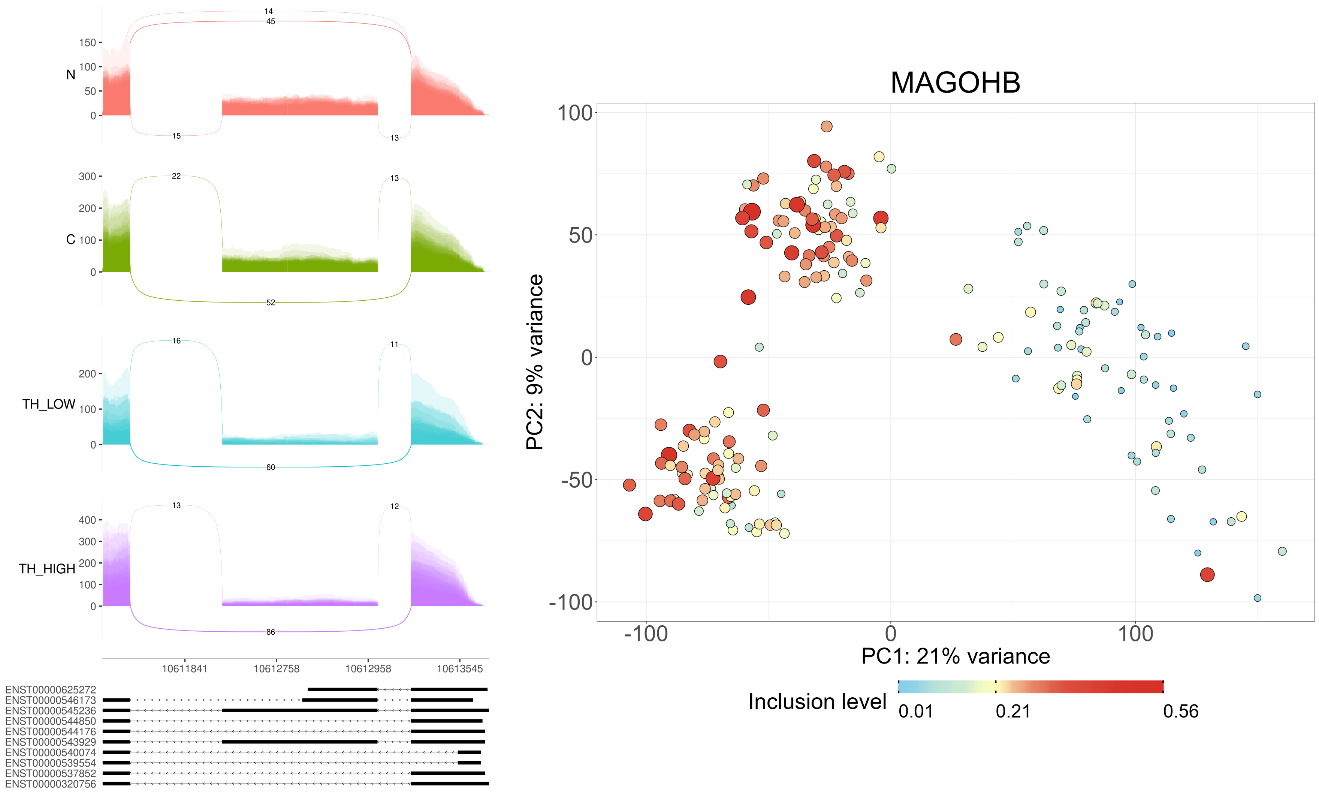

### Figure S13: Alternative splicing in the gene MAGOHB.

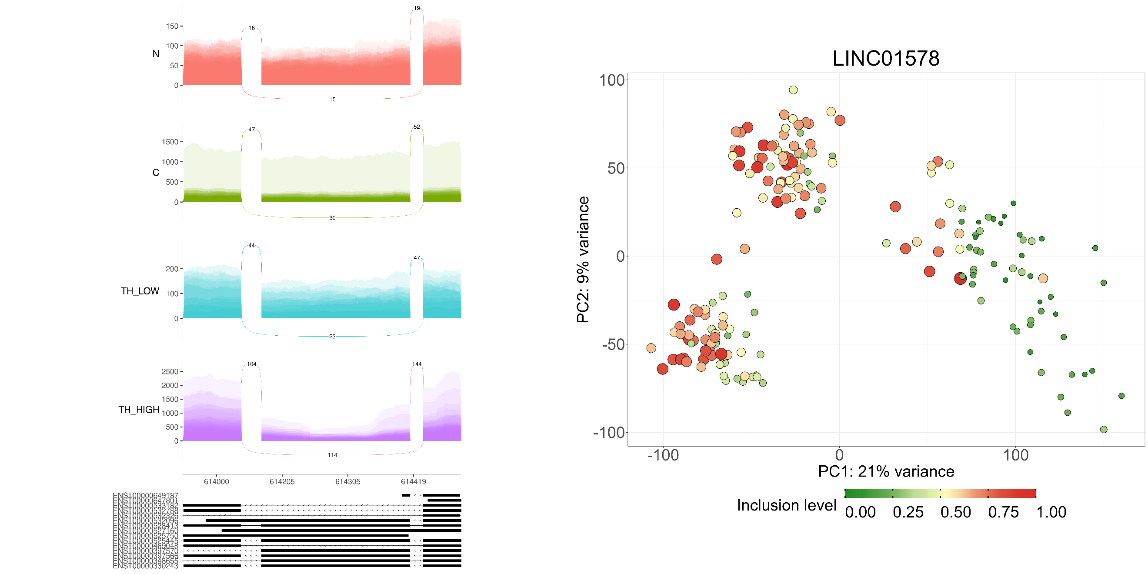
 Sashimi and feature plots of inclusion levels for a cassette exon in the gene MAGOHB.

This gene has multiple isoforms (Rehman, Chandra, and Singh 2021) and is involved in mRNA splicing (Barreiro et al. n.d.).

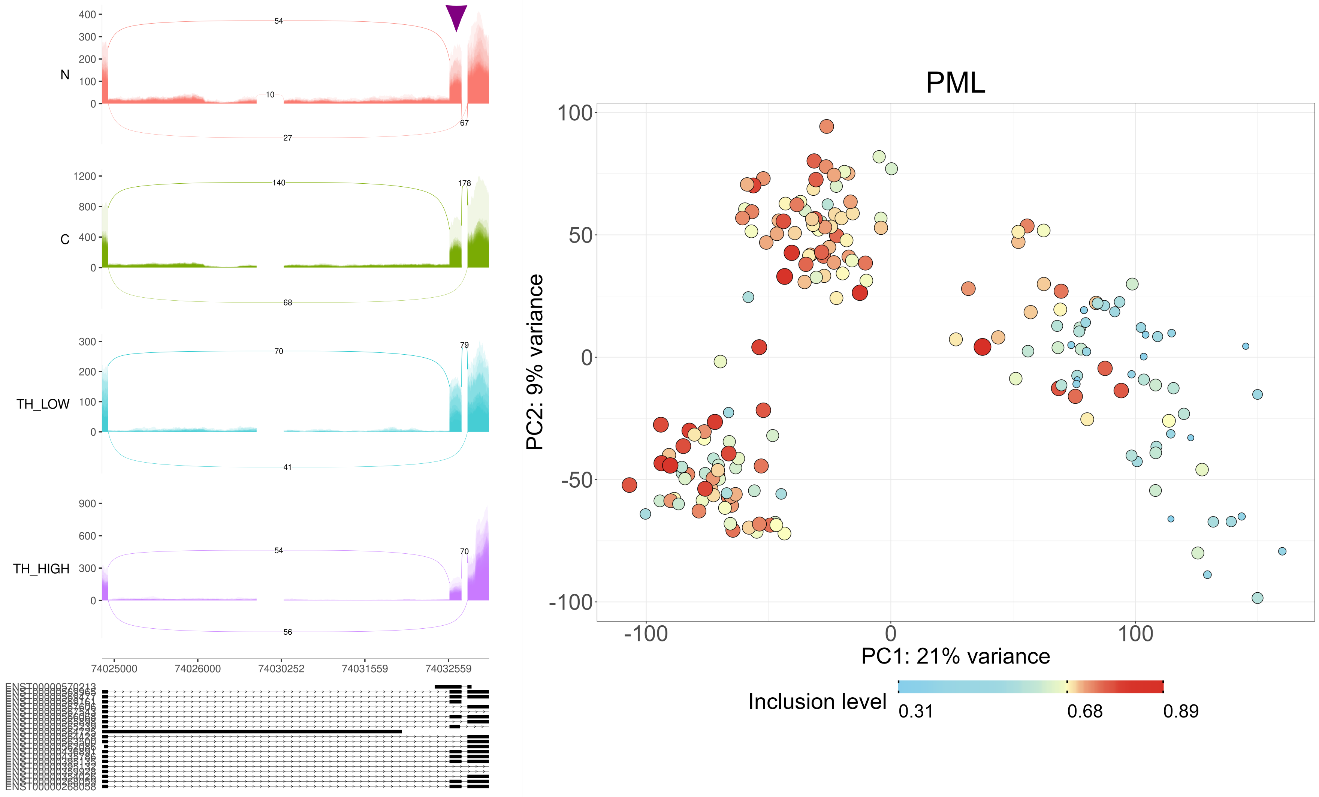

### Figure S14: Alternative splicing in the gene PML.

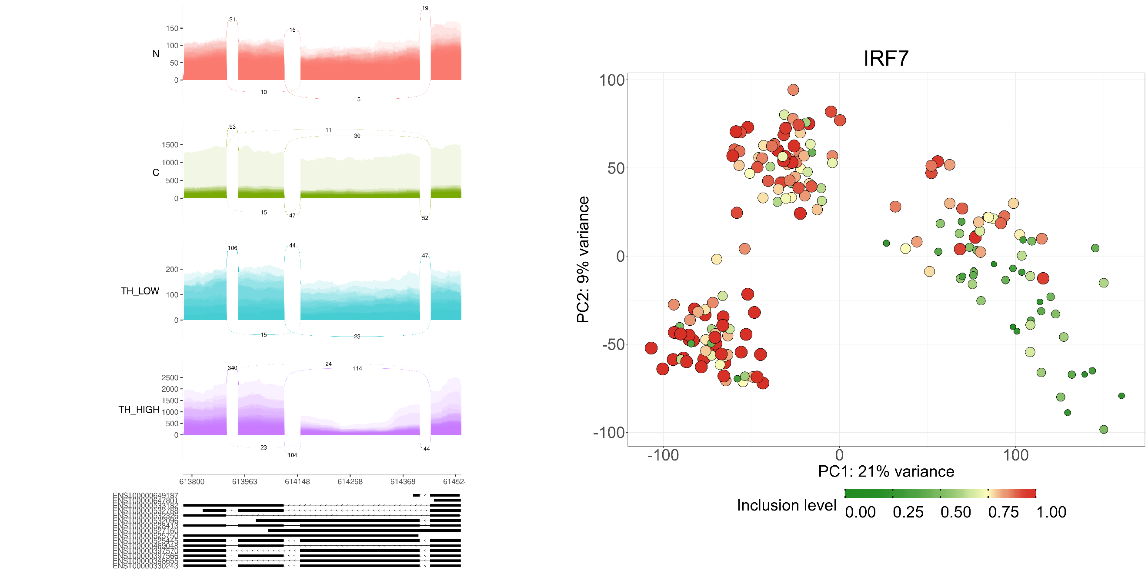

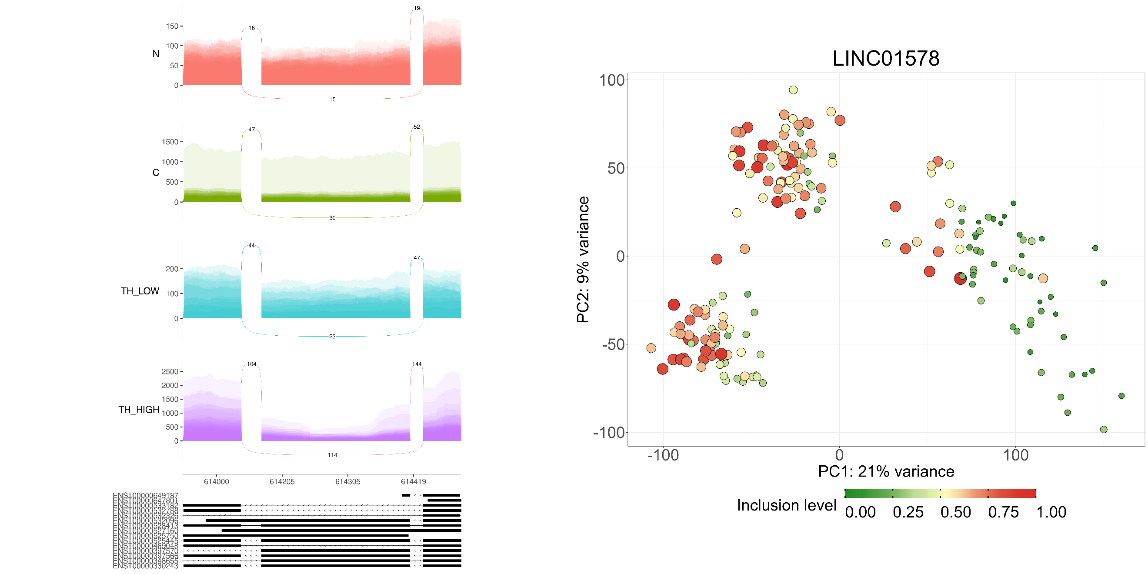
 Sashimi and feature plots of inclusion levels for cassette exon no.5 in the gene PML (Nisole et al. 2013).

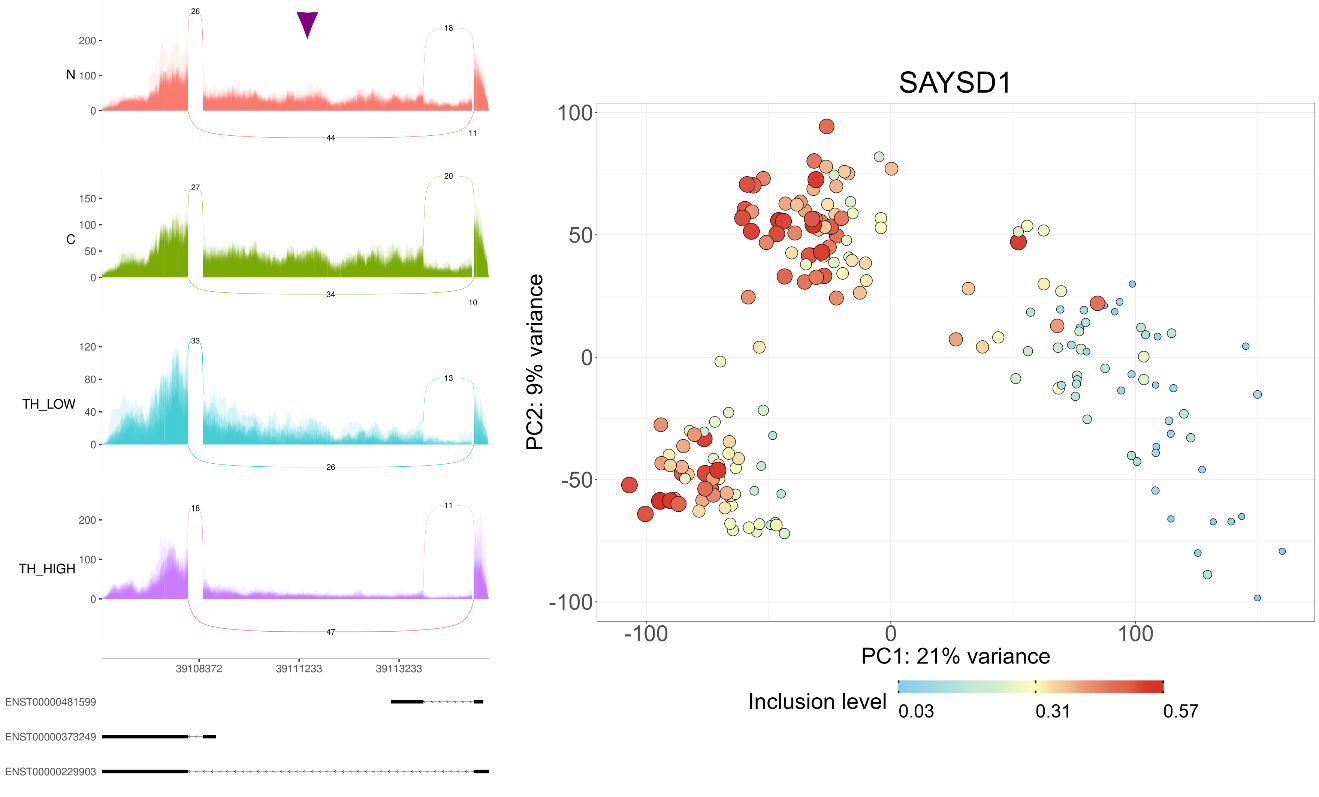

### Figure S15: Alternative splicing in the gene SAYSD1.

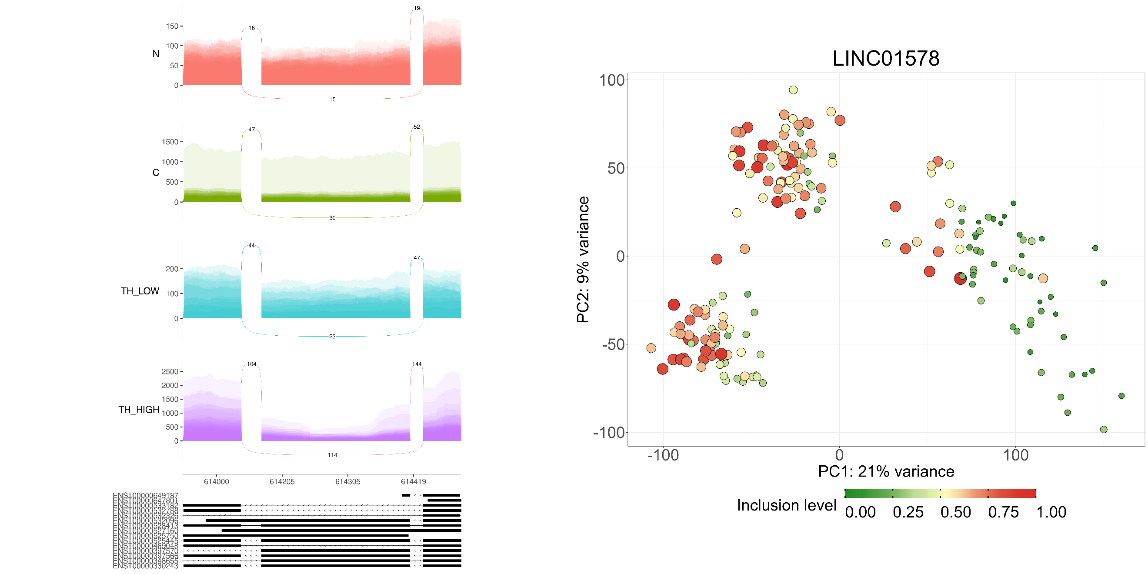
 Sashimi and feature plots of inclusion levels for a cassette exon in the gene SAYSD1.

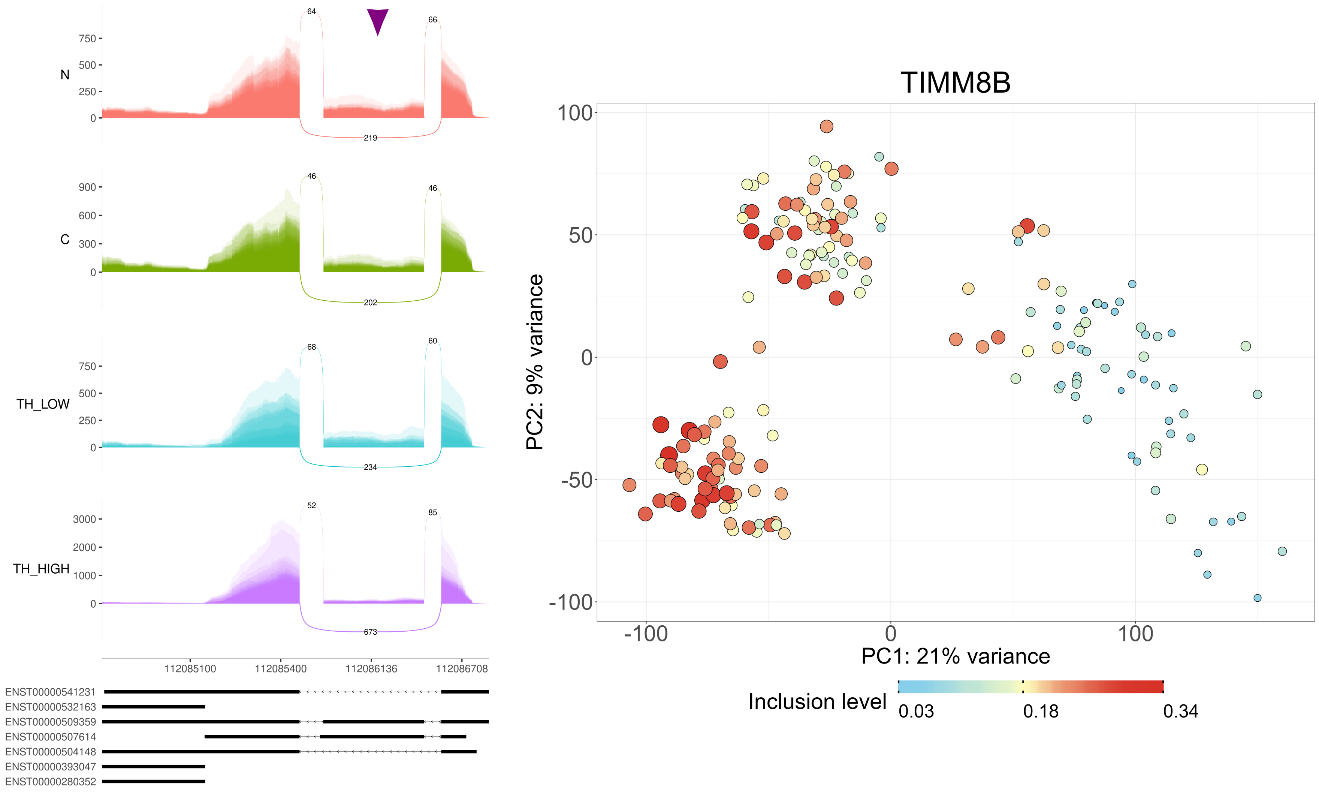

### Figure S16: Alternative splicing in the gene TIMM8B.

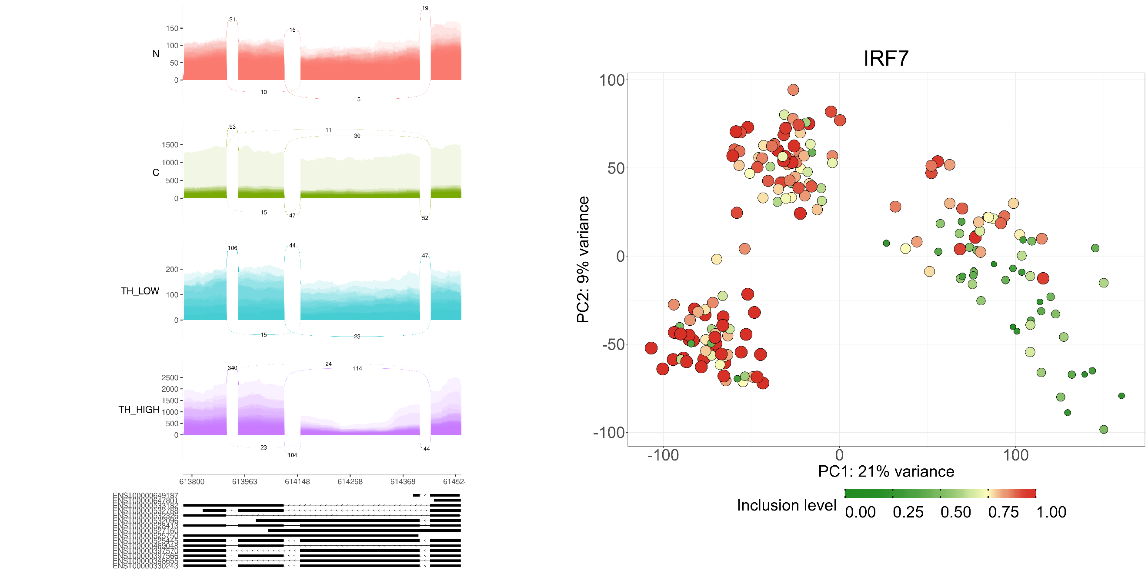

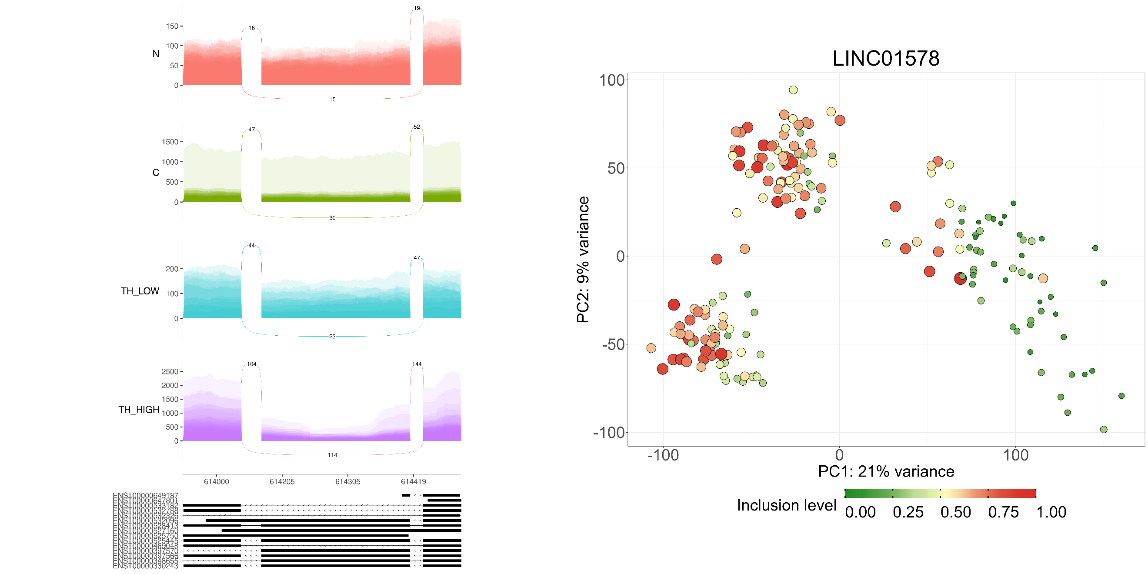
 Sashimi and feature plots of inclusion levels for a cassette exon in the gene TIMM8B.

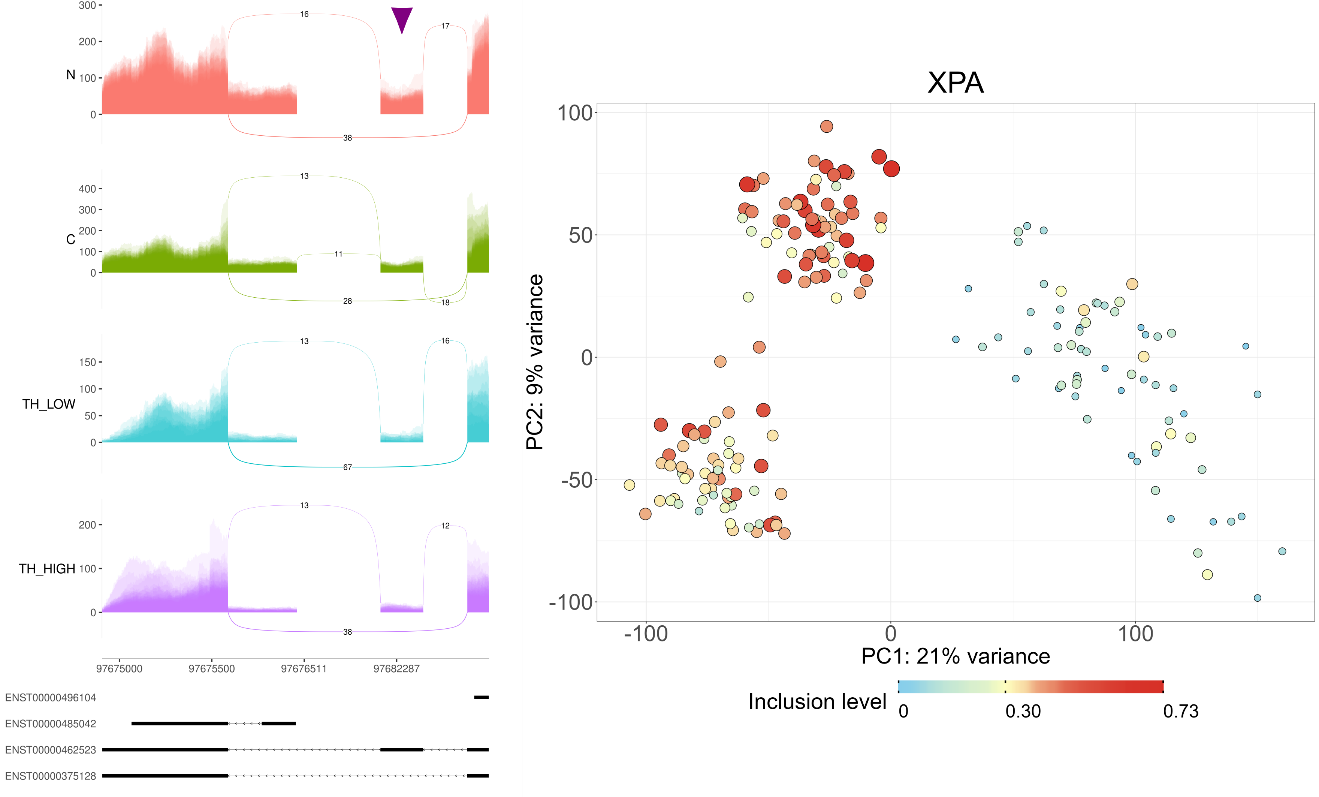

### Figure S17: Alternative splicing in the gene XPA.

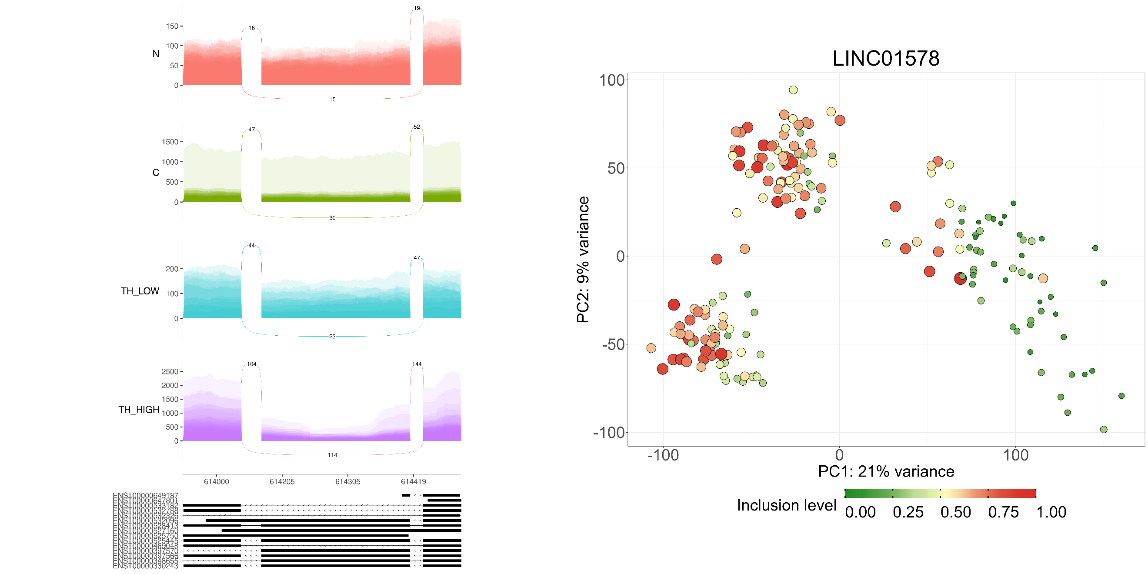
 Sashimi and feature plots of inclusion levels for a cassette exon in the gene XPA.

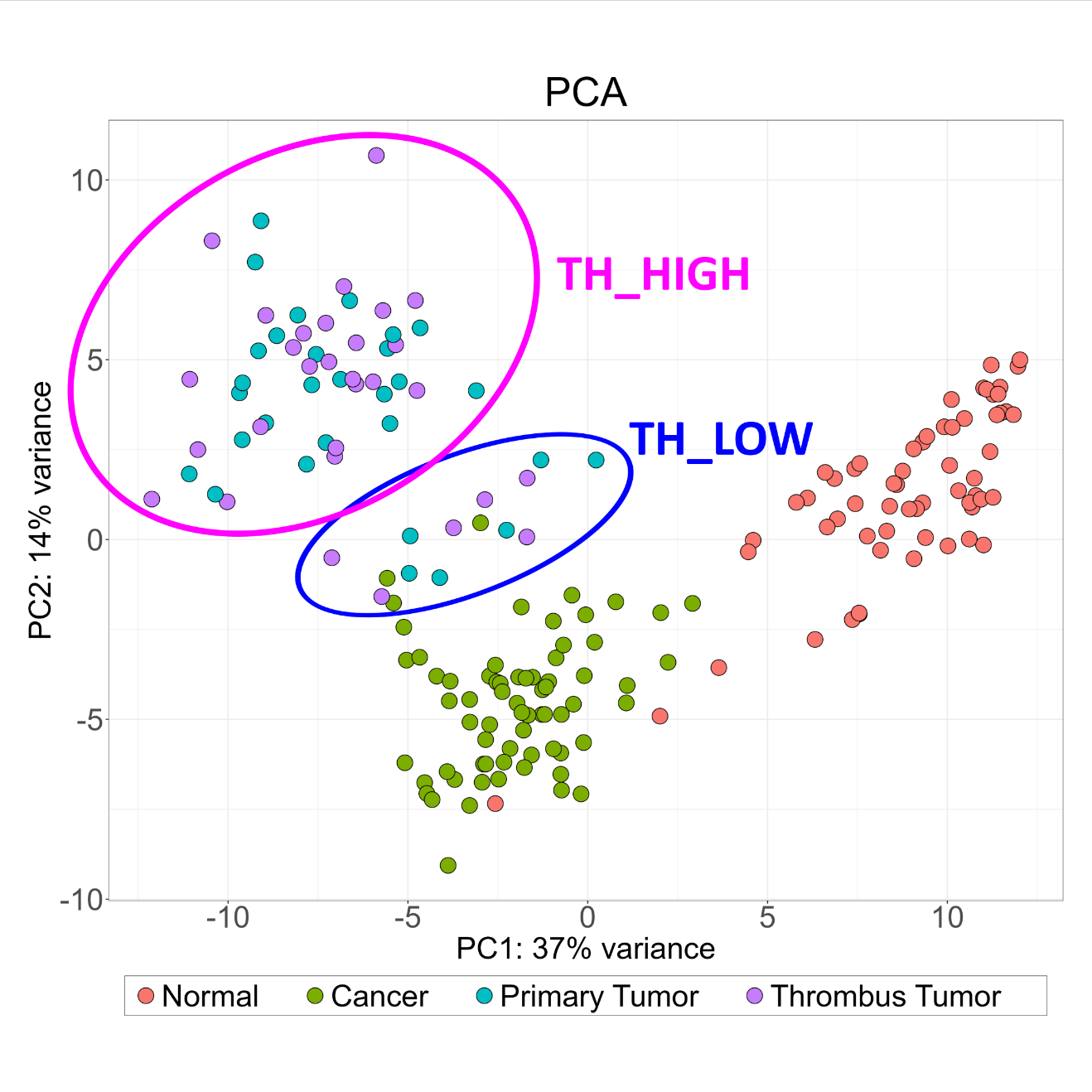

### Figure S18: A PCA plot of inclusion levels of cassette exons that were found to be significantly alternatively spliced in either of the three comparisons: N vs. C, C vs. PT&TT, and N vs. PT&TT (FDR=0 and |ΔPSI|>0.2).

It can be seen that a sub group of thormbi samples (PT&TT) is located in the proximity of the tumors without thrombi (C). We designated this subgroup as “TH_LOW” and all the other thrombi as “TH_HIGH”.

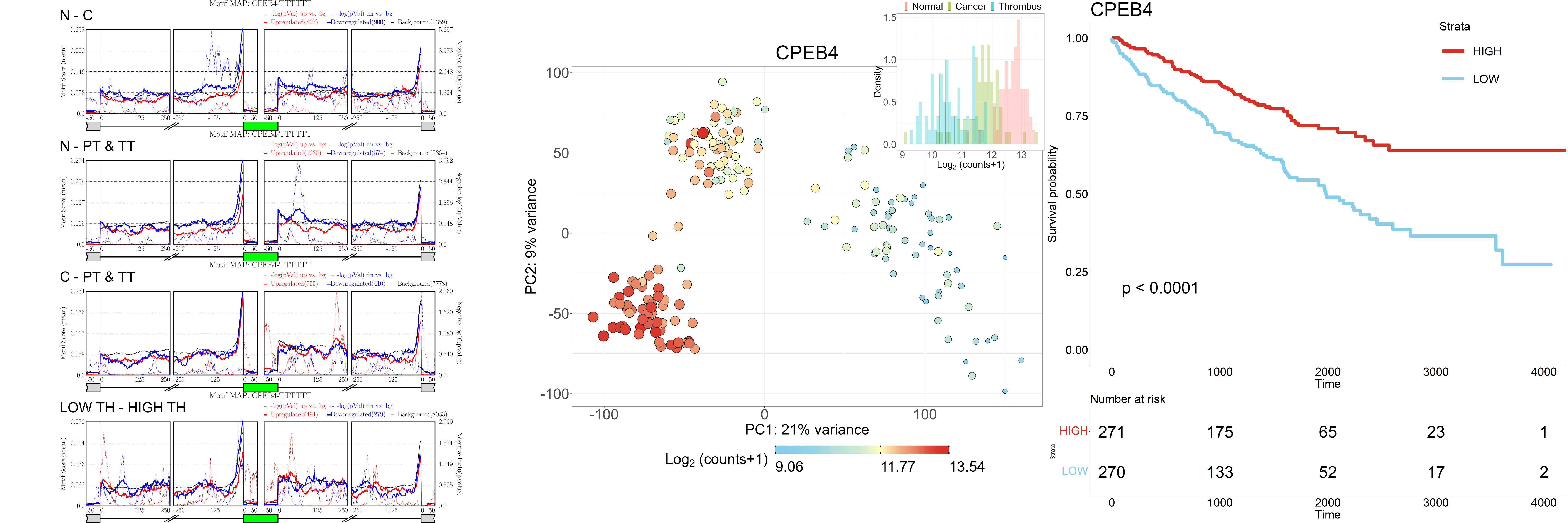

### Figure S19: CPEB4 is a putative splicing regulator associated with ccRCC progression.

Left panel shows enrichment plots for RNA binding motifs. Middle panel shows a feature plot for expression levels (red/large - high expression, blue/small – low expression). Right panel shows Kaplan-Meier curves based on the clear-cell RCC dataset (KIRC) from the TCGA database. Although rMAPS did not detect a significant motif enrichment, the expression levels and survival analysis of this gene indicate that it is associated with tumor progression.

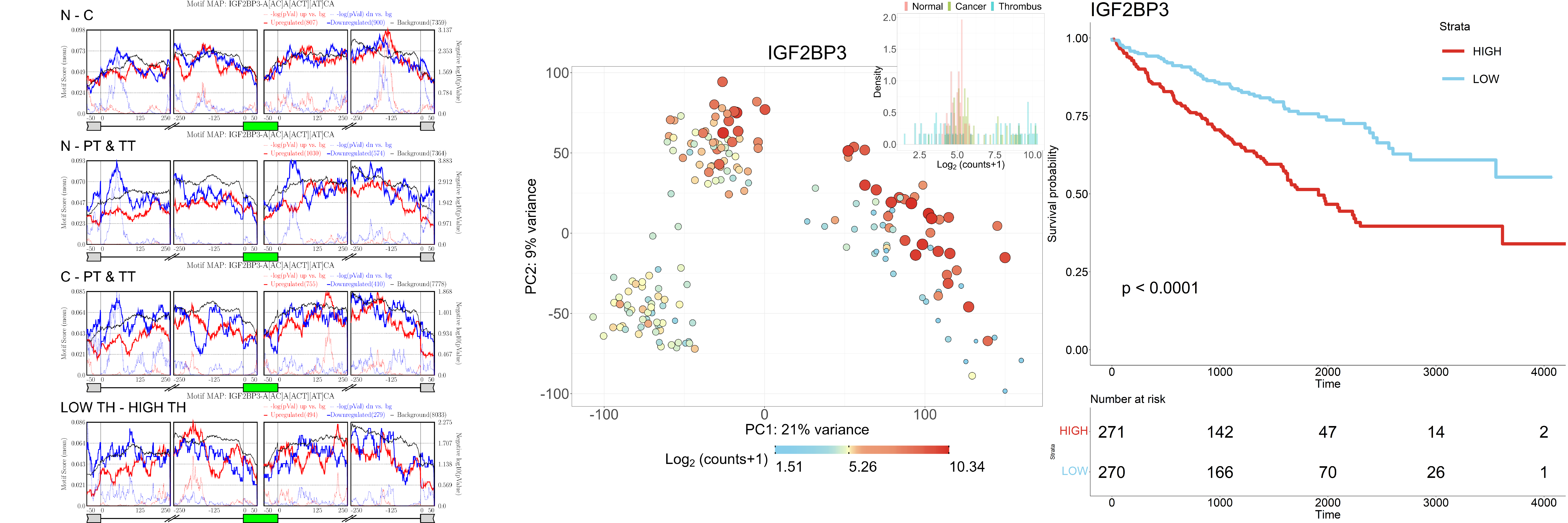

### Figure S20: IGF2BP3 is a putative splicing regulator associated with ccRCC progression.

Left panel shows enrichment plots for RNA binding motifs. Middle panel shows a feature plot for expression levels (red/large - high expression, blue/small – low expression). Right panel shows Kaplan-Meier curves based on the clear-cell RCC dataset (KIRC) from the TCGA database.

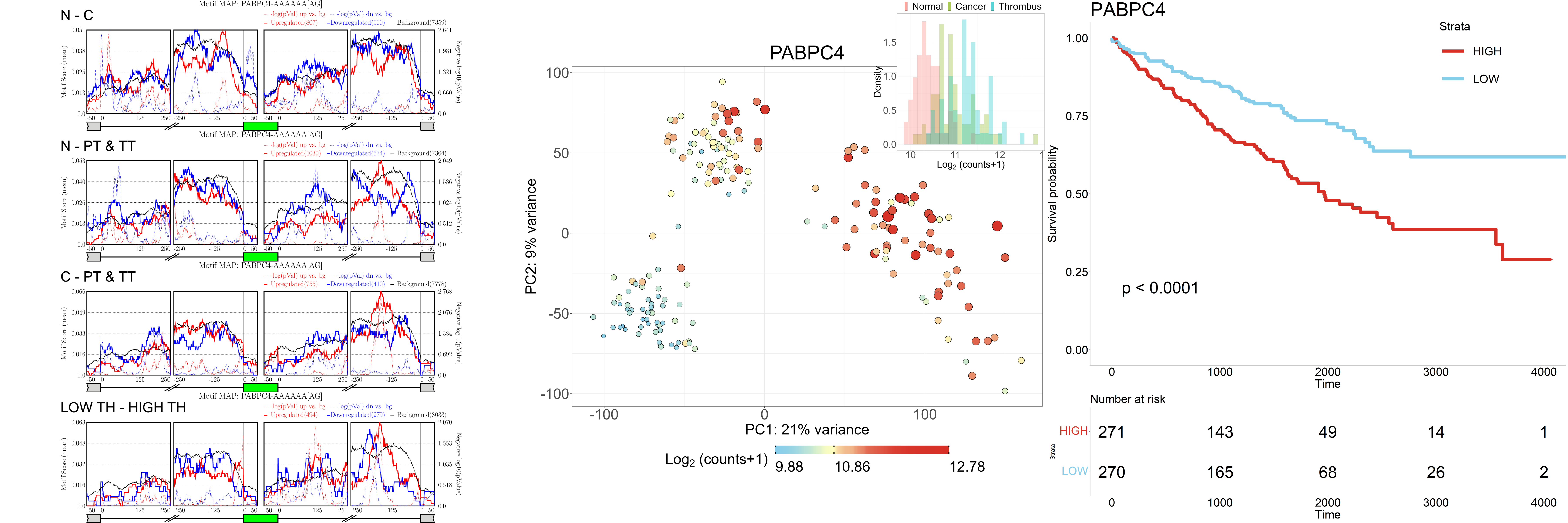

### Figure S21: PABPC4 is a putative splicing regulator associated with ccRCC progression.

Left panel shows enrichment plots for RNA binding motifs. Middle panel shows a feature plot for expression levels (red/large - high expression, blue/small – low expression). Right panel shows Kaplan-Meier curves based on the clear-cell RCC dataset (KIRC) from the TCGA database. Although rMAPS did not detect a significant motif enrichment, the expression levels and survival analysis of this gene indicate that it is associated with tumor progression.

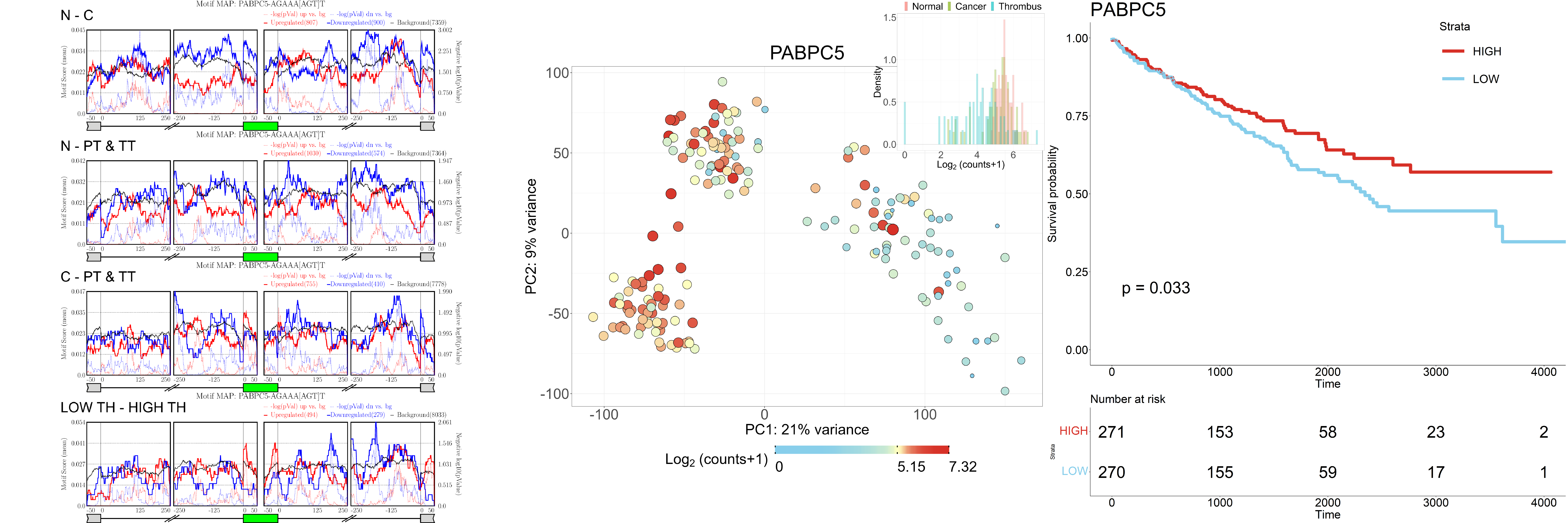

### Figure S22: PABPC5 is a putative splicing regulator associated with ccRCC progression.

Left panel shows enrichment plots for RNA binding motifs. Middle panel shows a feature plot for expression levels (red/large - high expression, blue/small – low expression). Right panel shows Kaplan-Meier curves based on the clear-cell RCC dataset (KIRC) from the TCGA database. Although rMAPS did not detect a significant motif enrichment, the expression levels and survival analysis of this gene indicate that it is associated with tumor progression.

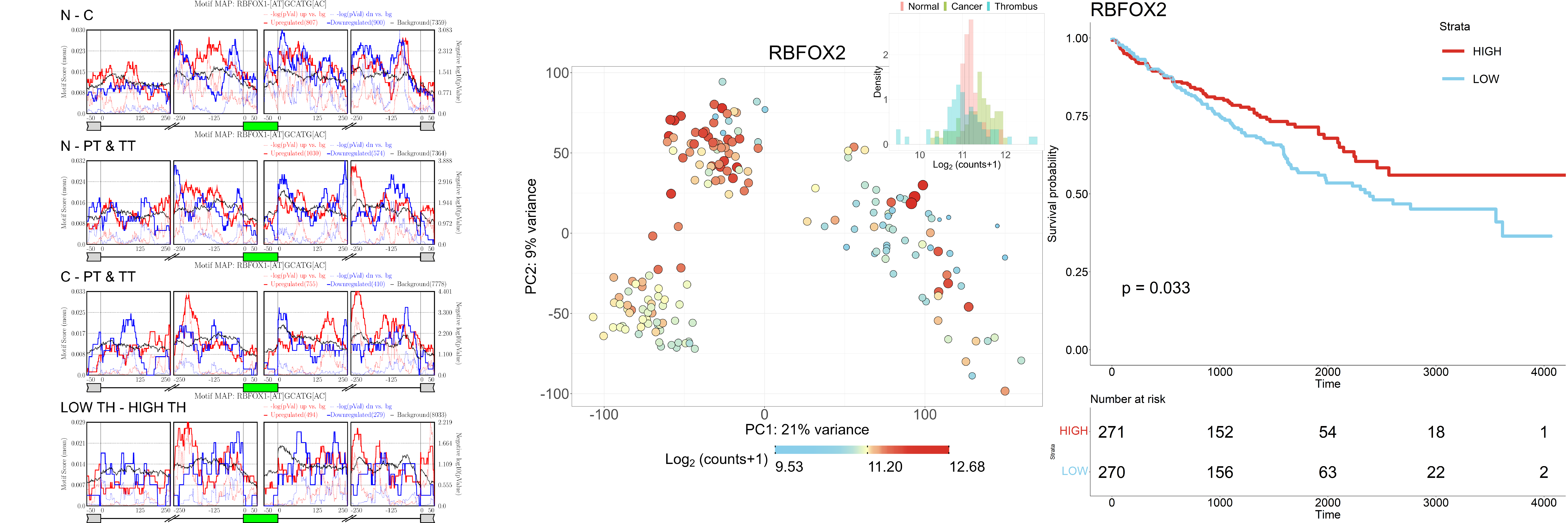

### Figure S23: RBFOX2 is a putative splicing regulator associated with ccRCC progression.

Left panel shows enrichment plots for RNA binding motifs. Middle panel shows a feature plot for expression levels (red/large - high expression, blue/small – low expression). Right panel shows Kaplan-Meier curves based on the clear-cell RCC dataset (KIRC) from the TCGA database. Although rMAPS did not detect a significant motif enrichment, the expression levels and survival analysis of this gene indicate that it is associated with tumor progression.

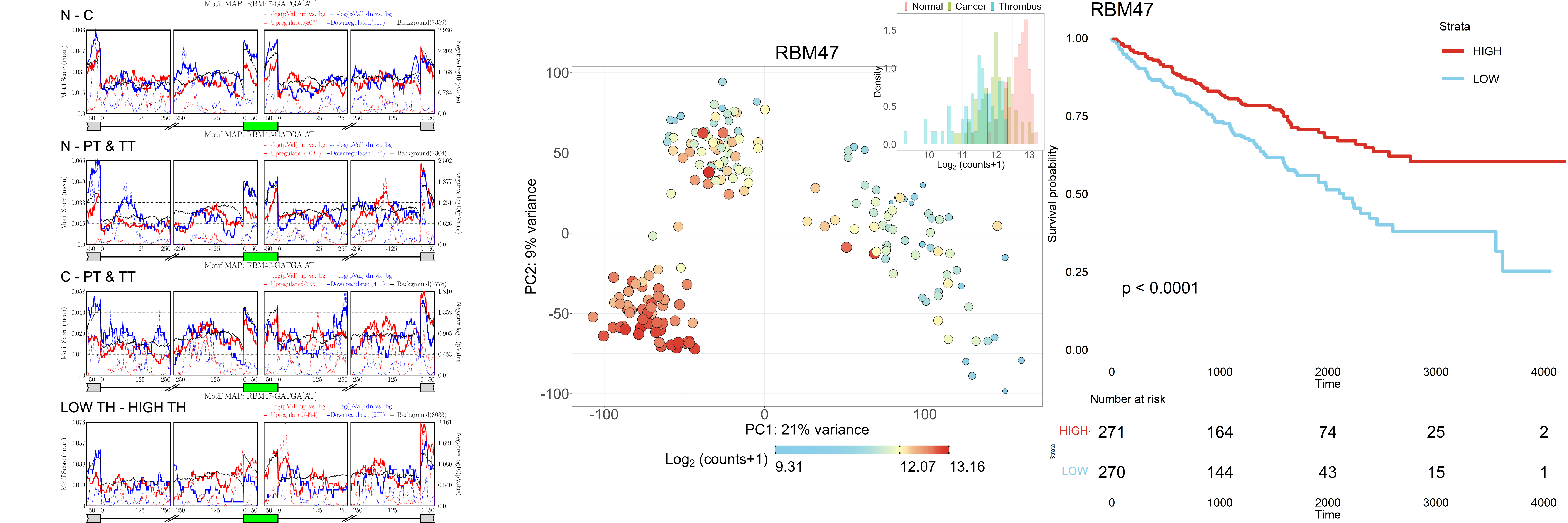

### Figure S24: RBM47 is a putative splicing regulator associated with ccRCC progression.

Left panel shows enrichment plots for RNA binding motifs. Middle panel shows a feature plot for expression levels (red/large - high expression, blue/small – low expression). Right panel shows Kaplan-Meier curves based on the clear-cell RCC dataset (KIRC) from the TCGA database. Although rMAPS did not detect a significant motif enrichment, the expression levels and survival analysis of this gene indicate that it is associated with tumor progression.

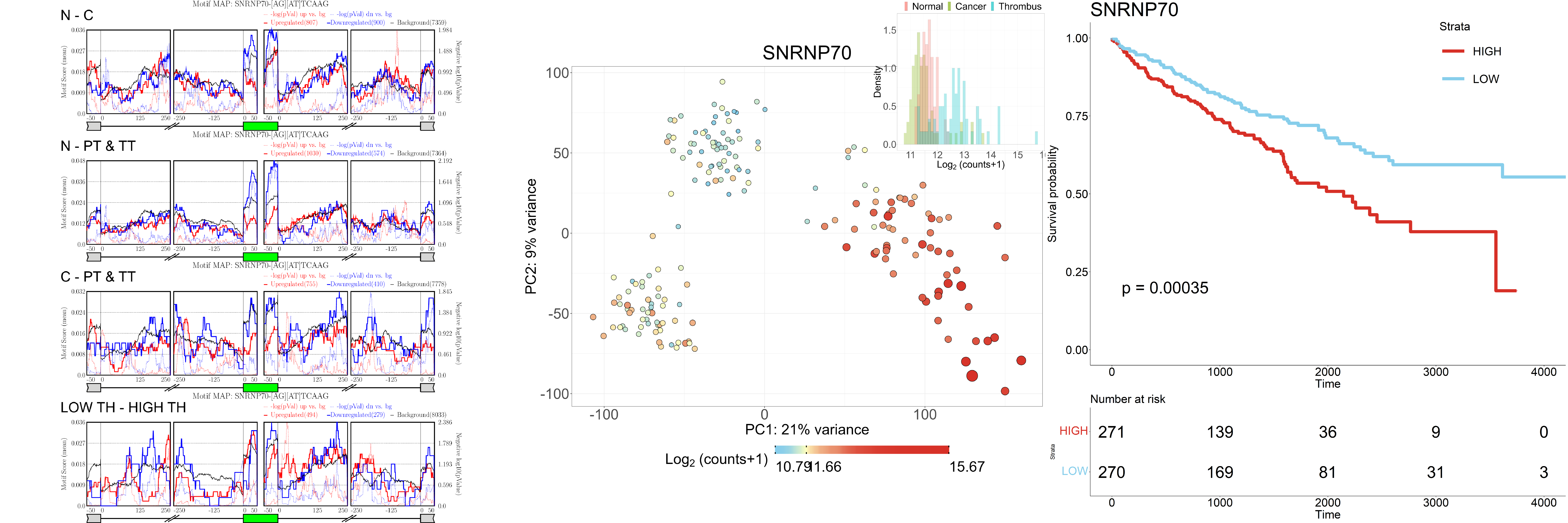

### Figure S25: SNRNP70 is a putative splicing regulator associated with ccRCC progression.

Left panel shows enrichment plots for RNA binding motifs. Middle panel shows a feature plot for expression levels (red/large - high expression, blue/small – low expression). Right panel shows Kaplan-Meier curves based on the clear-cell RCC dataset (KIRC) from the TCGA database. Although rMAPS did not detect a significant motif enrichment, the expression levels and survival analysis of this gene indicate that it is associated with tumor progression.

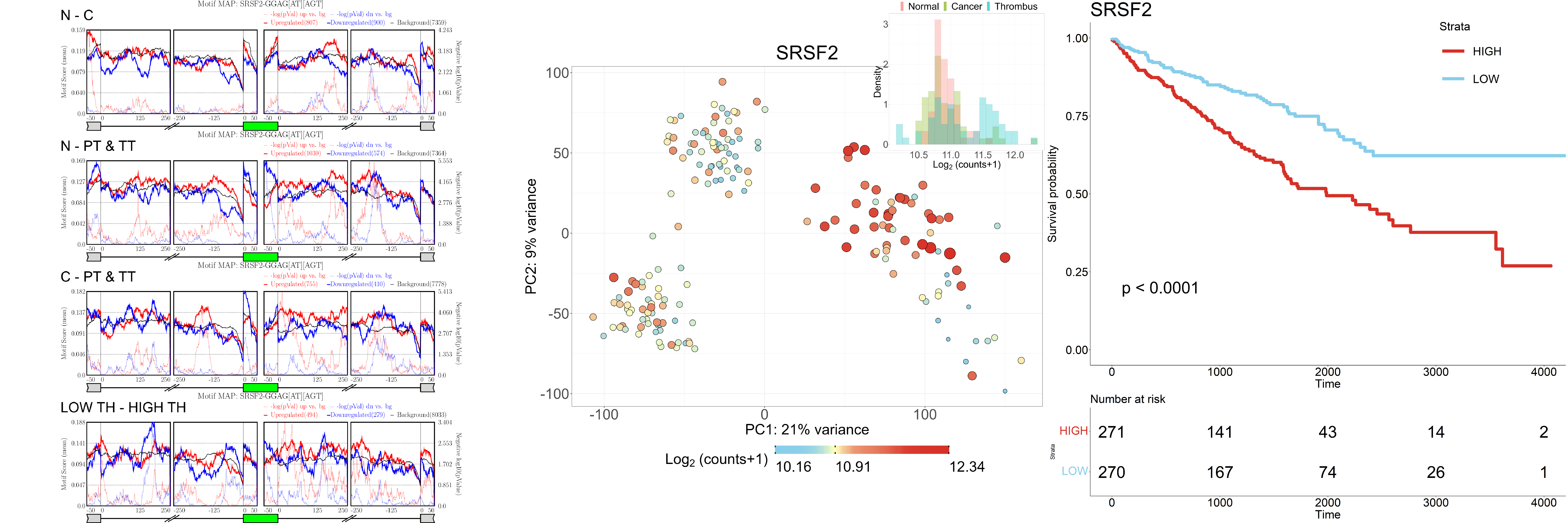

### Figure S26: SRSF2 is a putative splicing regulator associated with ccRCC progression.

Left panel shows enrichment plots for RNA binding motifs. Middle panel shows a feature plot for expression levels (red/large - high expression, blue/small – low expression). Right panel shows Kaplan-Meier curves based on the clear-cell RCC dataset (KIRC) from the TCGA database. Although rMAPS did not detect a significant motif enrichment, the expression levels and survival analysis of this gene indicate that it is associated with tumor progression.

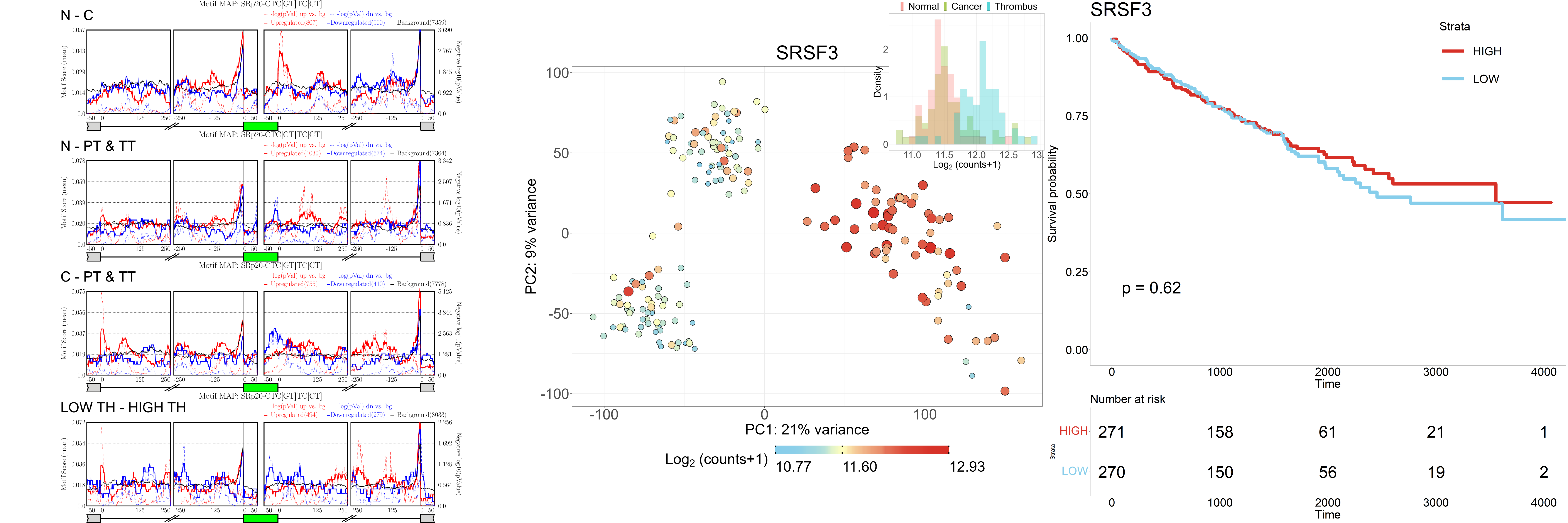

### Figure S27: SRSF3 is a putative splicing regulator associated with ccRCC progression.

Left panel shows enrichment plots for RNA binding motifs. Middle panel shows a feature plot for expression levels (red/large - high expression, blue/small – low expression). Right panel shows Kaplan-Meier curves based on the clear-cell RCC dataset (KIRC) from the TCGA database. Although survival analysis did not detect a significant clinical effect, the expression levels of this gene, and to some extent rMAPS motif enrichment as well, indicate that it is associated with tumor progression.

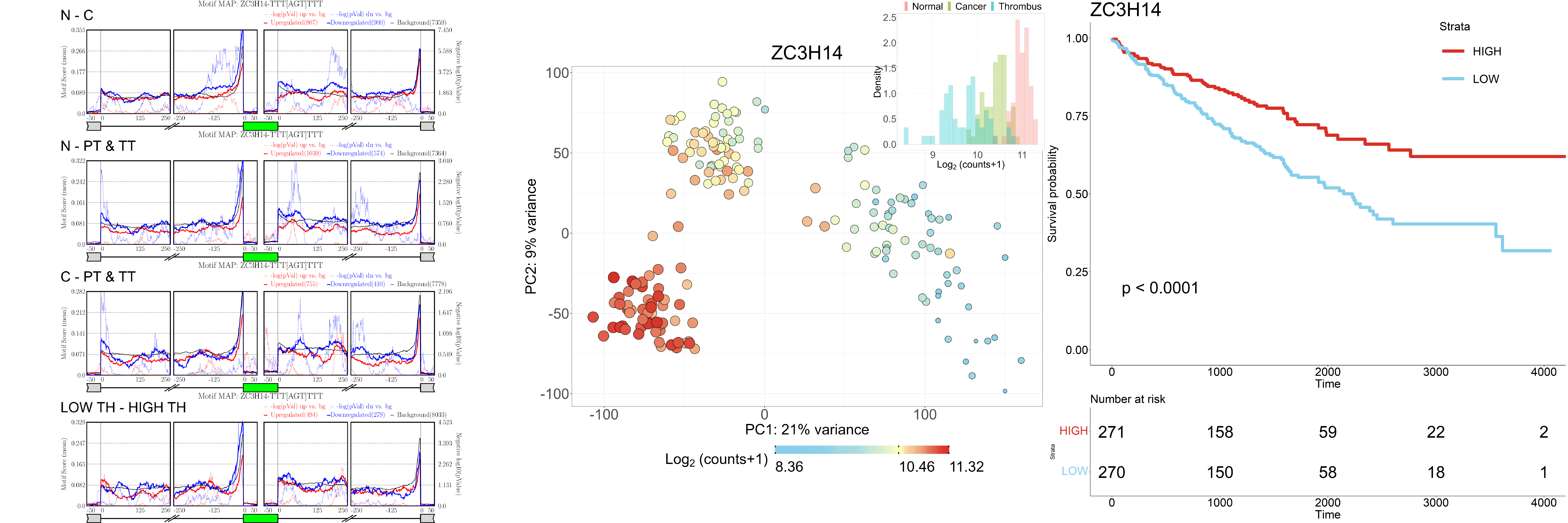

### Figure S28: ZC3H14 is a putative splicing regulator associated with ccRCC progression.

Left panel shows enrichment plots for RNA binding motifs. Middle panel shows a feature plot for expression levels (red/large - high expression, blue/small – low expression). Right panel shows Kaplan-Meier curves based on the clear-cell RCC dataset (KIRC) from the TCGA database. Although rMAPS did not detect a significant motif enrichment, the expression levels and survival analysis of this gene indicate that it is associated with tumor progression.

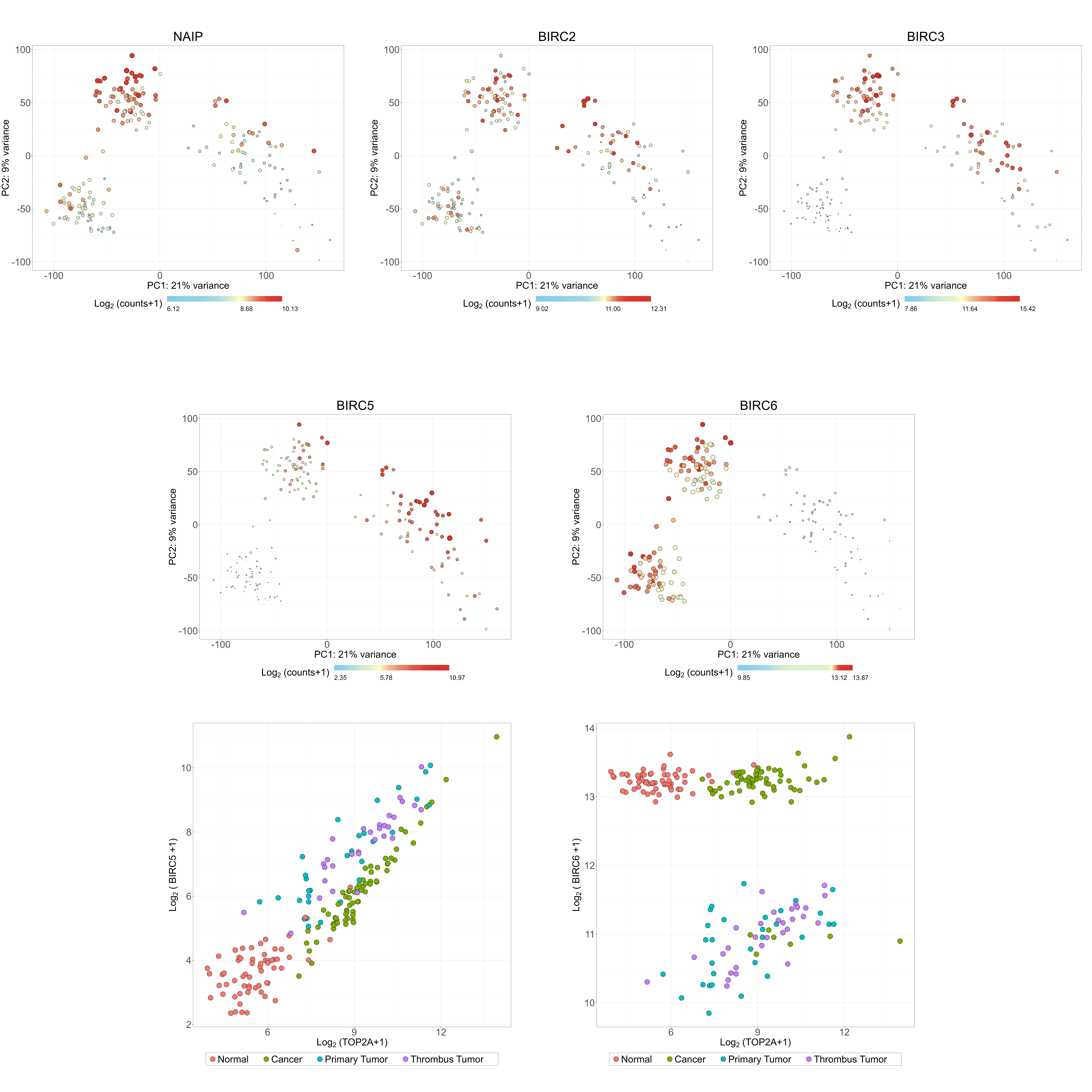

### Figure S29: Heterogeneous expression behavior of the BIRC gene family.

The genes BIRC1 (NAIP), BIRC2, and BIRC3, which are known to be associated with anti-apoptosis, are over-expressed in some, but not all tumors. The genes BIRC5 and BIRC6 are known to be associated with the cell cycle. BIRC5 is over-expressed in a subset of tumors and is strongly correlated to TOP2A.

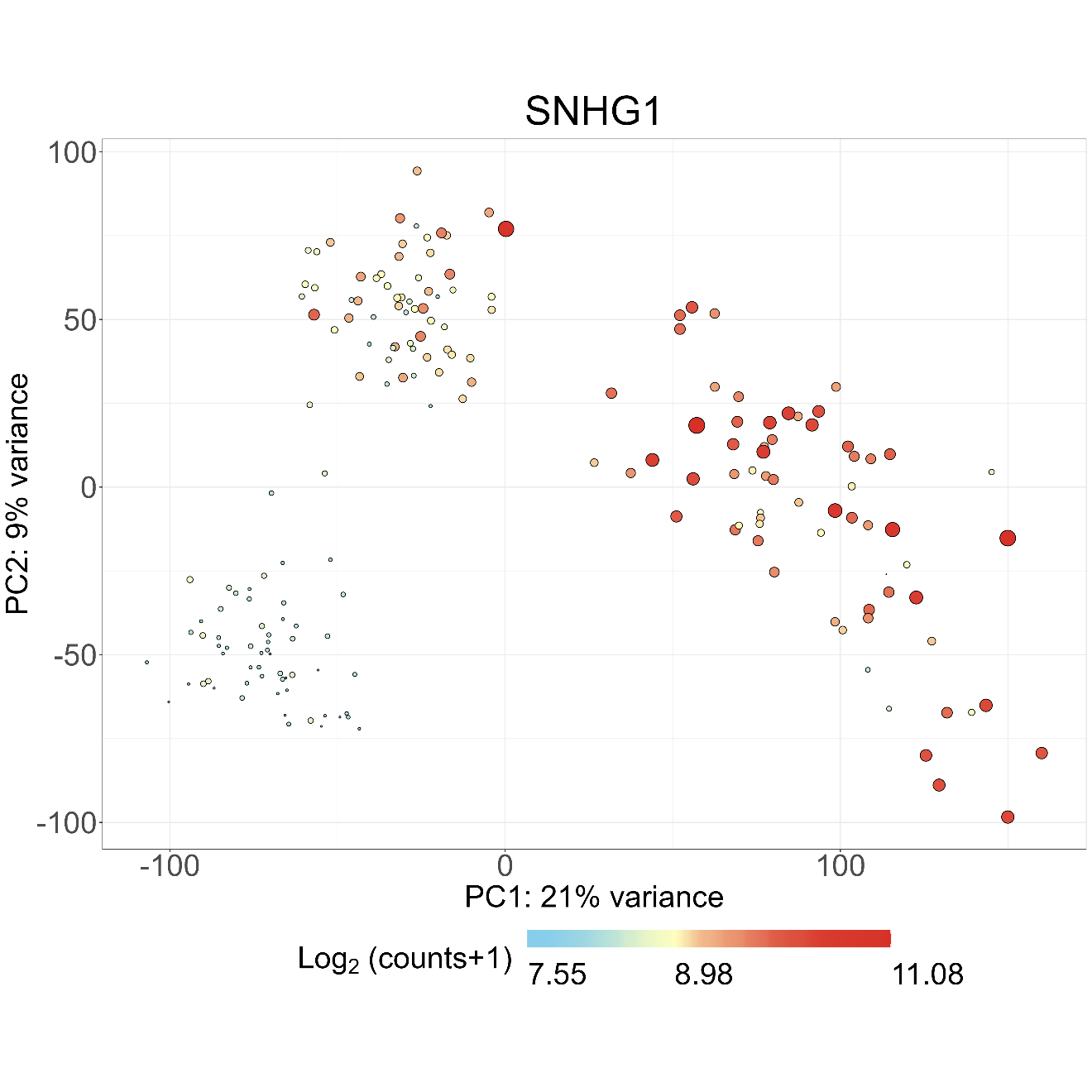

### **Figure S30: SNHG1 is over-expressed in cancer and further elevated in thrombi.**

Shown is a feature plot of SNHG1 (red/large - high expression, blue/small – low expression).
